# Single-cell transcriptomics and cell-specific proteomics reveals molecular signatures of sleep

**DOI:** 10.1101/2020.12.18.423331

**Authors:** Pawan K. Jha, Utham K. Valekunja, Sandipan Ray, Mathieu Nollet, Akhilesh B. Reddy

**Author notes:** Correspondence: Akhilesh B. Reddy. These authors contributed equally to the work (Pawan K. Jha, Utham K. Valekunja).

## Abstract

Every day, we sleep for a third of the day. Sleep is important for cognition, brain waste clearance, metabolism, and immune responses. The molecular mechanisms governing sleep are largely unknown. Here, we used a combination of single cell RNA sequencing and cell-type specific proteomics to interrogate the molecular underpinnings of sleep. Different cell types in three important brain regions for sleep (brainstem, cortex, and hypothalamus) exhibited diverse transcriptional responses to sleep need. Sleep restriction modulates astrocyte-neuron crosstalk and sleep need enhances expression of specific sets of transcription factors in different brain regions. In cortex, we also interrogated the proteome of two major cell types: astrocytes and neurons. Sleep deprivation differentially alters the expression of proteins in astrocytes and neurons. Similarly, phosphoproteomics revealed large shifts in cell-type specific protein phosphorylation. Our results indicate that sleep need regulates transcriptional, translational, and post-translational responses in a cell-specific manner.

## INTRODUCTION

Sleep is an essential component of daily life. The explicit function of sleep is still a mystery. Insufficient or disturbed sleep is associated with the accumulation of brain waste, cognitive impairments, increased risk of metabolic abnormalities, and suppressed immune responses (Besedovsky et al., 2019; Palmer and Alfano, 2017; Reutrakul and Cauter, 2018; Xie et al., 2013). Restoration of many of these abnormalities by sleep suggests that prolonged wakefulness imposes a load on the system that is alleviated by sleep (Benington and Heller, 1995; Maquet, 1995; Tononi and Cirelli, 2014; Xie et al., 2013). The progressive rise of sleep need during wakefulness, and its dissipation during sleep, maintain sleep homeostasis (Borbely, 1982; Borbély et al., 2016).

Sleep homeostasis is an intrinsic process that regulates sleep need by the cumulative and interactive action of multiple brain regions and cells (Terao et al., 2008; Thompson et al., 2010; Vyazovskiy and Harris, 2013). Sleep-wake cycles are regulated mainly in the hypothalamus, brainstem, and cortex of the brain (Rosenwasser, 2009; Saper and Fuller, 2017). Although prior work has described bulk transcriptional responses of various brain regions following acute sleep restriction (Cirelli et al., 2004; Elliott et al., 2014; Noya et al., 2019; Terao et al., 2006; Winrow et al., 2009), cell-type specific changes in gene and protein expression have not been delineated. Thus, there is a substantial gap in our knowledge concerning the cell types that are responsible for transcriptional changes during sleep deprivation, and whether different cell types are affected in different ways by sleep deprivation. Here, we set out to plug this gap by determining the molecular changes within individual brain cells across sleep-wake states.

We used single cell RNA sequencing (scRNA-seq) to compare the cellular composition and transcriptomic profiles of brainstem, cortex and hypothalamus of mice going through different sleep treatments. For all of the major cell populations that are responsive to the sleep treatments, we provide a comprehensive census of gene expression and analysis of molecular pathways affected. We show that sleep need regulates the transcriptional profiles of brain cells in the three brain regions differently. Given that astrocyte-neuron signaling plays a crucial role in homeostatic brain function (Mederos et al., 2018), we delineated ligand-receptor interactions that are altered by sleep treatments within these cell types. Our scRNA-seq data enabled us to identify uniquely enriched transcription factors controlling changes in gene expression in the brain cells from sleep deprived animals. We also analyzed the change in protein levels, and post-translational phosphorylation, following sleep treatments in cortical astrocytes and neurons. Taken together, this study significantly advances our understanding of the molecular changes in sleep homeostasis at the individual brain cell level and serves as a foundation for further cell-specific interrogation of sleep need.

## RESULTS

### Sleep phenotyping and experimental design

We sleep restricted wild-type C57BL/6J mice by prolonging their wakefulness into the light phase of the light-dark cycle, from ZT0 to ZT12 (ZT; zeitgeber time). This is when mice normally obtain 60-70% of their daily sleep (Tang and Sanford, 2002). As expected, the mice showed increased sleep need, as evidenced by elevated slow wave activity (SWA) and delta power (1-4 Hz) in their electroencephalography (EEG) traces in the recovery period after sleep deprivation (Borbely, 1982; Borbély et al., 2016) (Figures 1A and S1). In addition, animals spent significantly more time asleep following sleep restriction compared to their corresponding baseline hours (Figure S1A). This indicates accumulated sleep pressure during prolonged wakefulness.

**Figure 1.**
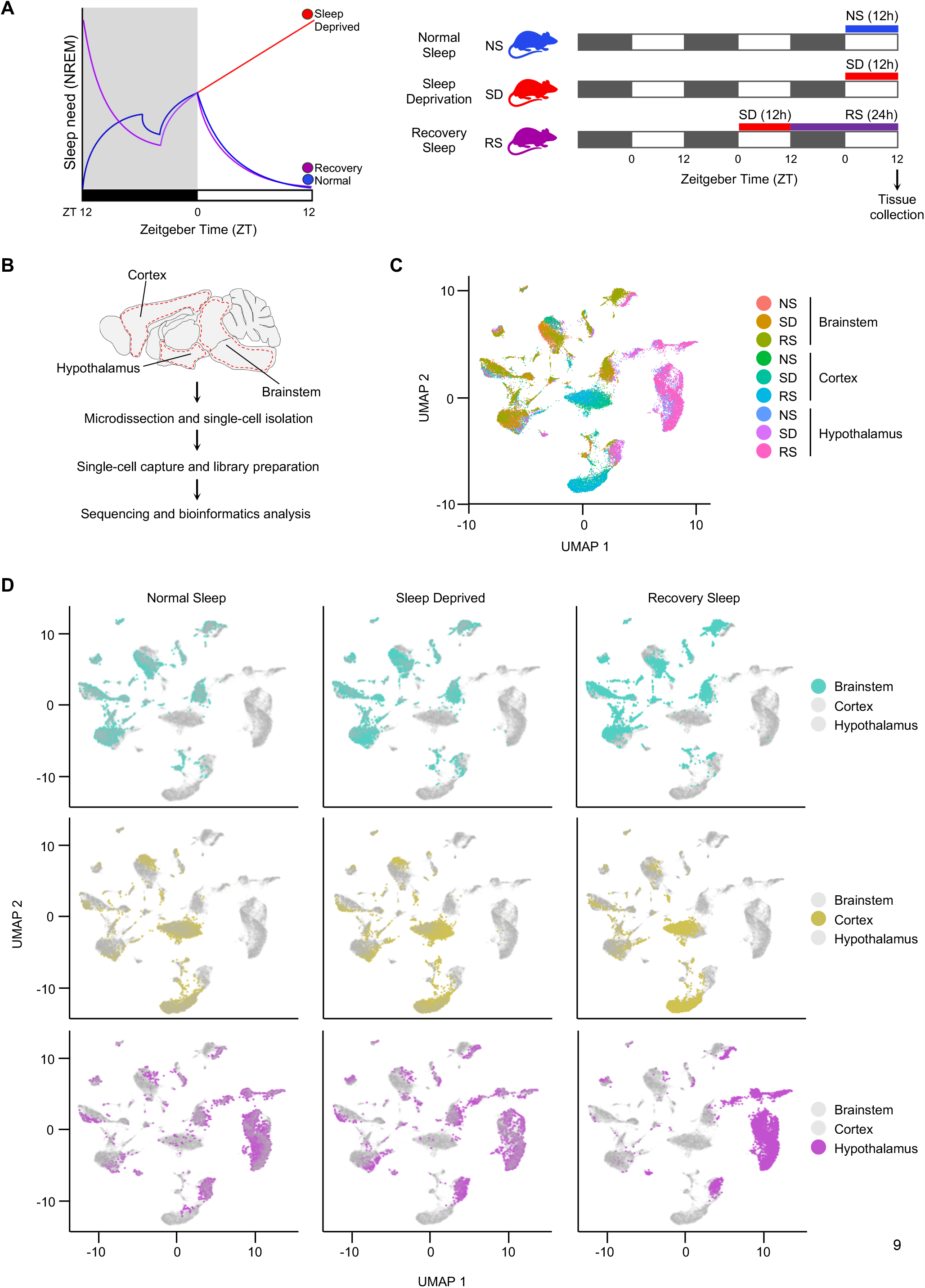
Transcriptional profile of brainstem, cortex and hypothalamic cells across sleep-wake states. (A) Sleep need model and experimental design (black bars represent darkness, white bars indicate periods of light). (B) Workflow for single-cell RNA-seq of cells from brainstem, cortex and hypothalamus, from microdissection to bioinformatics analysis. Dotted red line indicates areas of each brain region dissected for single cell sequencing. (C and D) UMAP visualization of 29,051 cells. Cells from Normal Sleep (NS), 12h Sleep Deprivation (SD) and 12h Sleep Deprivation followed by 24 h Recovery Sleep (RS) treatment groups from all three brain regions are clustered together (C) and separately highlighted (D). See also Figure S1 and S2.

We next divided the mice into three experimental groups: a control normal sleep (NS) group, a 12 h sleep deprived (SD) group, and a sleep deprivation followed by 24 h of recovery sleep (RS) group (Figure 1A). From each group, we collected samples from three brain regions (brainstem, cortex, and hypothalamus) that are pivotal for regulating sleep-wake cycles (Saper and Fuller, 2017; Saper et al., 2005). Importantly, we controlled for variation due to time of day by harvesting brain tissue at the same circadian phase (ZT12) (Figure 1A). We then performed single cell RNA sequencing (scRNA-seq) on cell suspensions from each brain region, in each sleep treatment group (Figure 1B).

### Identification and transcriptional profiling of brain cells across sleep-wake states

To understand how the change of sleep-wake states affects gene expression patterns within the cells of all three brain areas, we transcriptionally profiled cells and visualized them using a nonlinear dimensionality-reduction method, uniform manifold approximation and projection (UMAP) (Figure 1C and 1D) (Liu, 2020). Considering all three experimental sleep groups and the three brain regions together, we generated clusters from the quality criteria passed 29,051 cells (Figure S2A-S2C). When we analyzed each brain region individually, we found that individual cells were distributed among distinct clusters for each brain region, and there was no variation in these clusters for different sleep treatments (Figure 1D). This indicates there was no global shift in the molecular identity of cells in each brain region during sleep deprivation and recovery, which was as expected for terminally differentiated cells in the brain.

We next used UMAP to subdivide the globally defined clusters within the brain regions and used these subclusters to identify cell types based on known markers derived from existing single cell data, and the available literature (Chen et al., 2017; Mickelsen et al., 2019; Saunders et al., 2018; Zeisel et al., 2018) (Figures 2A-B, 3A-B, 4A-B, S2D-F, and Table S2). We classified 10,521 cells into 18 clusters within the brainstem, 8,451 cells into 14 clusters within the cortex, and 10,081 cells into 19 clusters within the hypothalamus. Cell types included astrocytes, endothelial cells, ependymocytes (EPC), hemoglobin-expressing vascular cells (HbVC), macrophages, microglia, neurons, oligodendrocytes (Oligo), pericytes and vascular smooth muscle cells, arterial (VSMCA) (Figures S2D-I).

**Figure 2.**
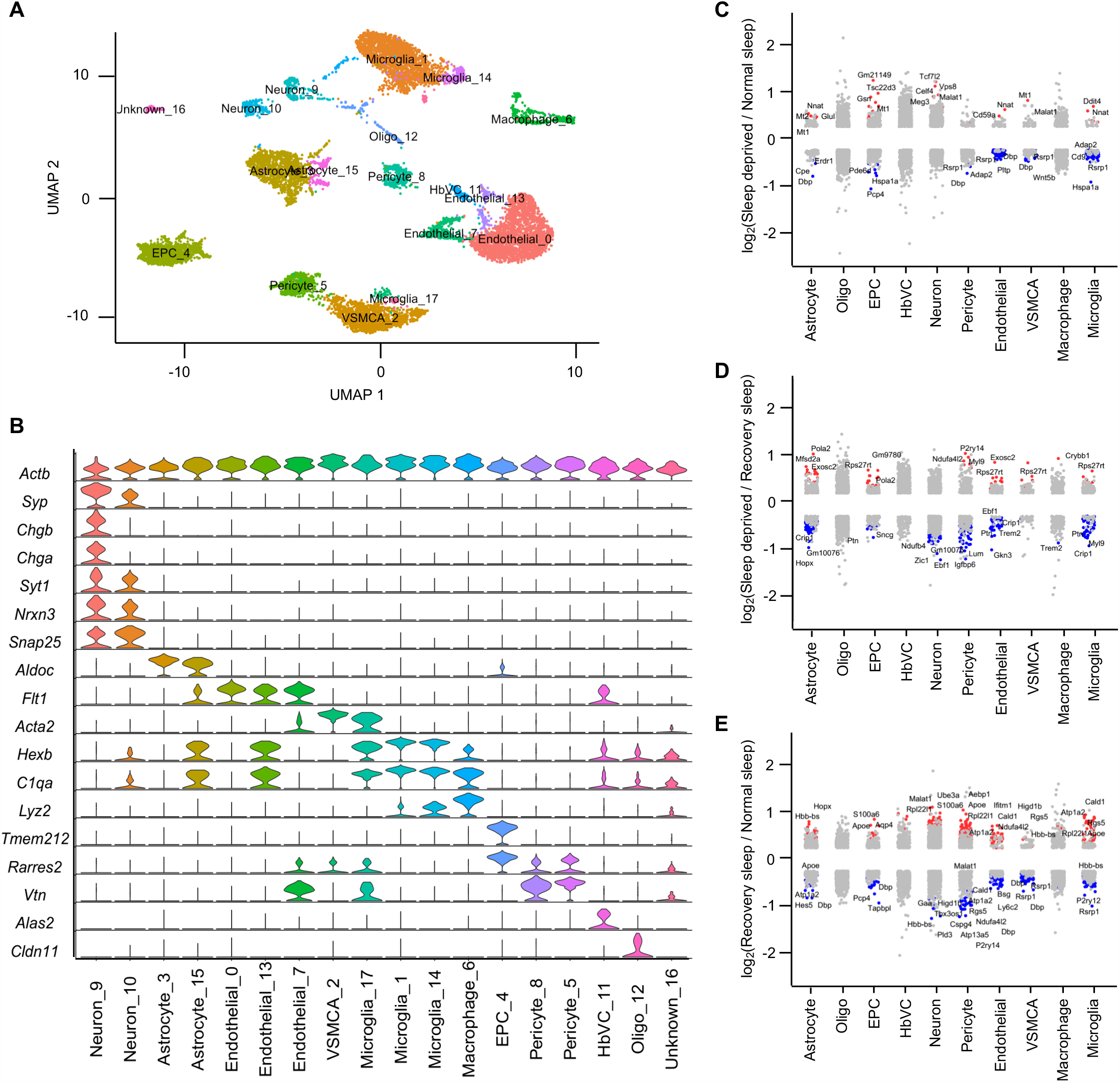
Sleep treatments alter the expression profile of cells in brainstem. (A) UMAP visualization of brainstem cells clustered. Cell clusters were colour coded and annotated based on their transcriptional profiles (see STAR Methods). (B) Violin plots for each cluster showing the expression level of selected known cell-type enriched markers. *Actb* (beta actin) is shown as a positive control in all cell types. (C-E) Strip chart showing changes in gene expression of Sleep Deprivation (SD)/Normal Sleep (NS) (C), Sleep Deprivation (SD)/Recovery Sleep (RS) (D), and (E) Recovery Sleep (RS)/Normal Sleep (NS) comparisons. Wilcoxon rank sum tests followed by false discovery rate (FDR) analysis were used to compare the groups. Significantly upregulated and downregulated genes are colour coded with red and blue, respectively (Bonferroni adjusted *P*-value < 0.1). Genes in grey are not significantly changed after sleep deprivation. See also Figures S2, S3 and S4, and Tables S2 and S3.

Next, to understand how the change in sleep need affects transcriptional profiles of cells in brainstem, cortex, and hypothalamus, we performed differential gene expression analysis of the three sleep treatment groups for each cell type we identified (Figures 2C-E, 3C-E, and 4C-E). We found distinct patterns of transcriptional response to sleep need in the astrocytes, neurons, endothelial cells, and microglia of all three brain regions. Pericytes, VSMCAs and EPCs of brainstem and hypothalamus also showed similar changes. Interestingly, compared to neurons, astrocytes showed larger alterations in brainstem and hypothalamus (Table S3). Of note, there was not a large change in gene expression patterns of Oligo, HbVC and macrophage cell types following sleep treatments (Figures 2C-E, 3C-E, and 4C-E). We then determined the overlap of sleep related alterations between previous bulk cell RNA-seq studies and our scRNA-seq data to assign cell types for those genes (Diessler et al., 2018; Gerstner et al., 2016; Scarpa et al., 2018). We were thus able to assign some expression changes found previously in bulk RNA profiling to astrocytes, neurons, endothelial cells, and microglia in cortex and hypothalamus using our novel single cell data (Figure S7A-B, Table S4).

**Figure 3.**
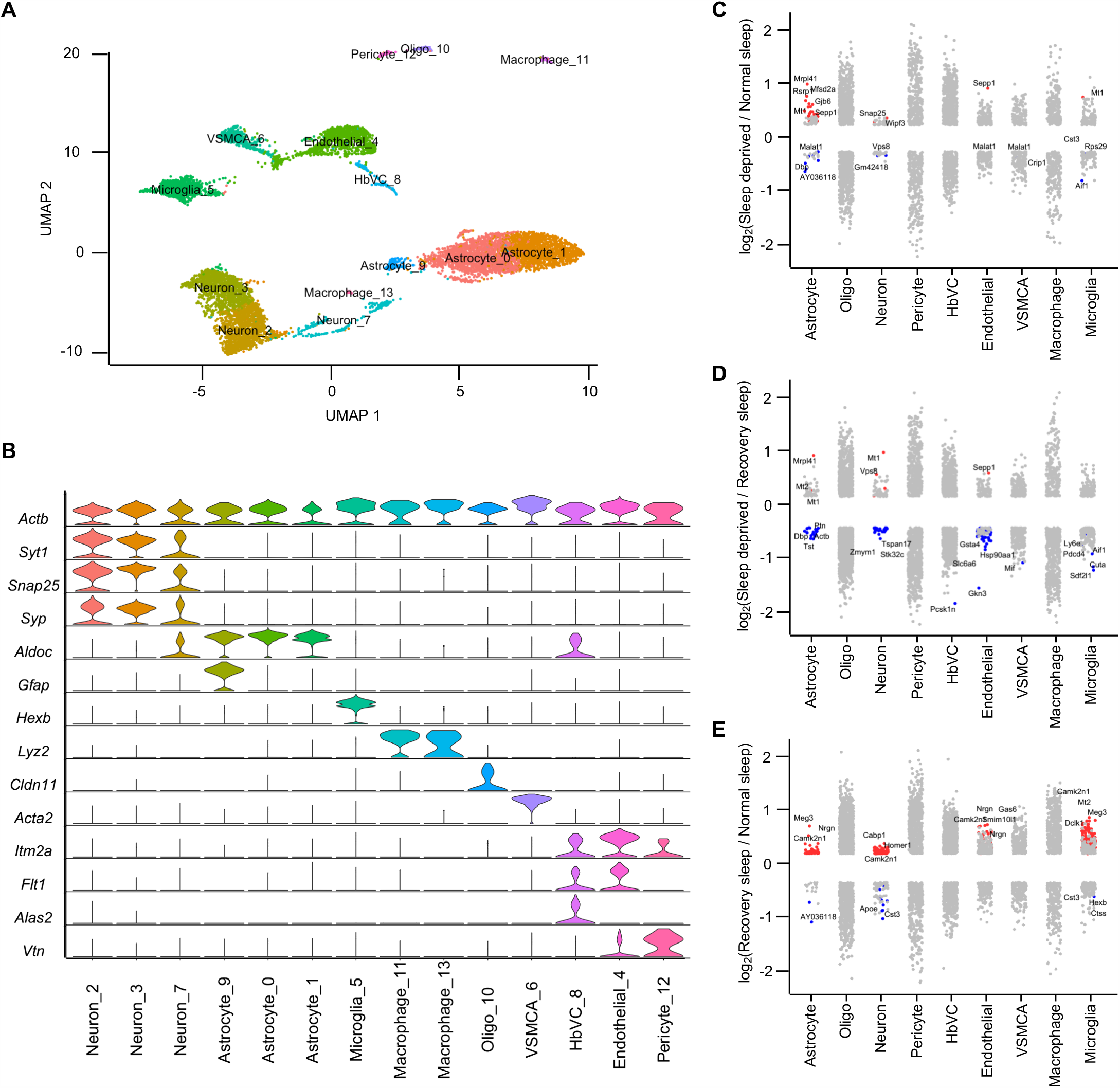
Sleep treatments alter the expression profile of cells in cortex. (A) UMAP visualization of brainstem cells clustered. Cell clusters were colour coded and annotated based on their transcriptional profiles (see STAR Methods). (B) Violin plots for each cluster showing the expression level of selected known cell-type enriched markers. *Actb* (beta actin) is shown as a positive control in all cell types. (C-E) Strip chart showing changes in gene expression of Sleep Deprivation (SD)/Normal Sleep (NS) (C), Sleep Deprivation (SD)/Recovery Sleep (RS) (D), and (E) Recovery Sleep (RS)/Normal Sleep (NS) comparisons. Wilcoxon rank sum tests followed by false discovery rate (FDR) analysis were used to compare the groups. Significantly upregulated and downregulated genes are colour coded with red and blue, respectively (Bonferroni adjusted *P*-value < 0.1). Genes in grey are not significantly changed after sleep deprivation. See also Figures S2, S3, S4 and S7, and Tables S2, S3, and S4.

**Figure 4.**
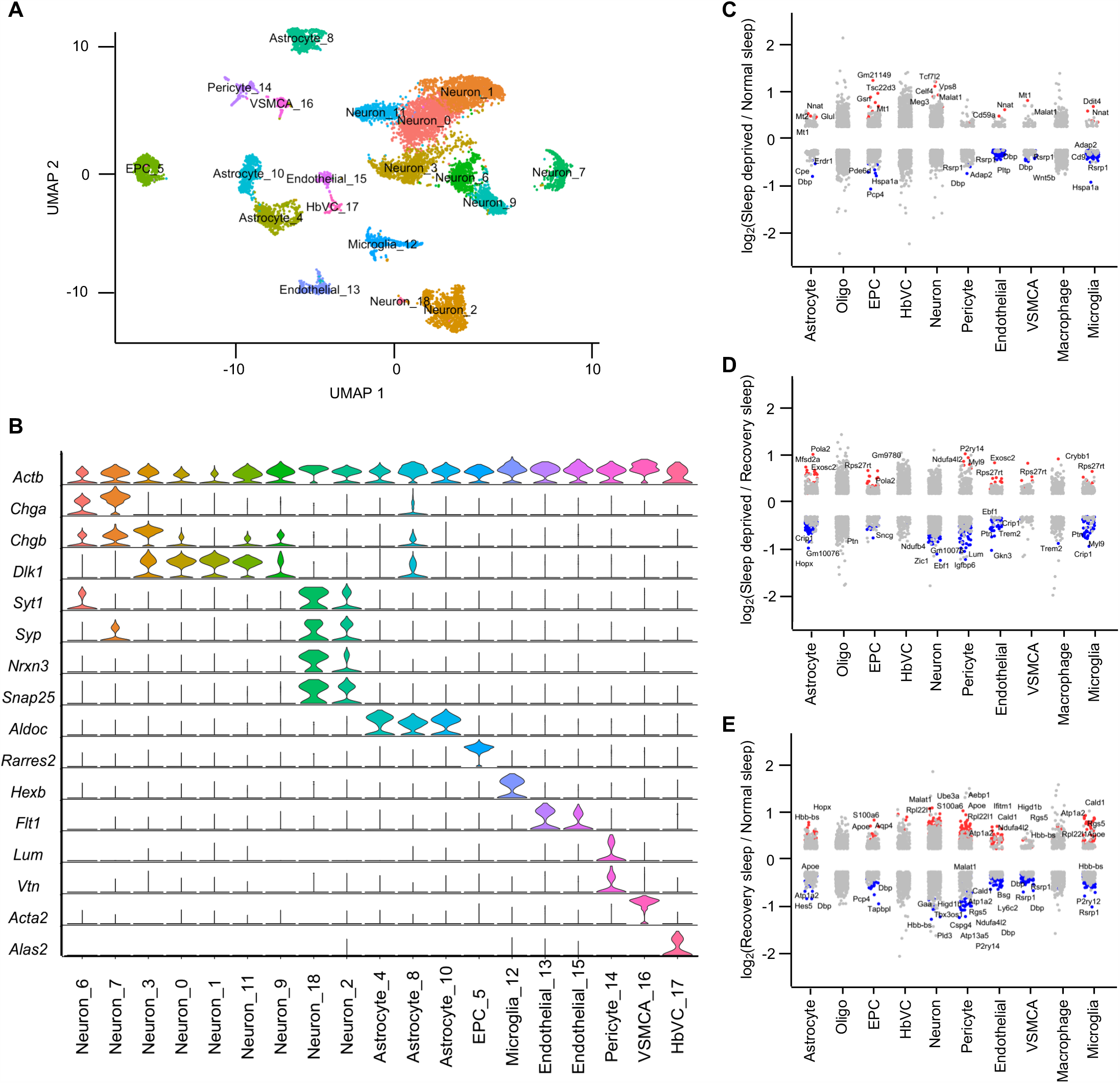
Sleep treatments alter the expression profile of cells in hypothalamus. (A) UMAP visualization of brainstem cells clustered. Cell clusters were colour coded and annotated based on their transcriptional profiles (see STAR Methods). (B) Violin plots for each cluster showing the expression level of selected known cell-type enriched markers. *Actb* (beta actin) is shown as a positive control in all cell types. (C-E) Strip chart showing changes in gene expression of Sleep Deprivation (SD)/Normal Sleep (NS) (C), Sleep Deprivation (SD)/Recovery Sleep (RS) (D), and (E) Recovery Sleep (RS)/Normal Sleep (NS) comparisons. Wilcoxon rank sum tests followed by false discovery rate (FDR) analysis were used to compare the groups. Significantly upregulated and downregulated genes are colour coded with red and blue, respectively (Bonferroni adjusted *P*-value < 0.1). Genes in grey are not significantly changed after sleep deprivation. See also Figures S2, S3, S4 and S7, and Tables S2, S3, and S4.

We next performed gene-annotation enrichment analysis of genes differentially expressed across sleep treatments in the cells of brainstem, cortex, and hypothalamus (Figures S3 and S4). In astrocytes, we found significant alterations in genes involved in ribosomal assembly, ribosomal RNA (rRNA) processing, and transmembrane transporter activity. In contrast, vesicle mediated transport in synapses and nucleotide/nucleoside phosphate metabolism were altered in neurons (Figures S3A-B, and S4A-B).

Cell types in different brain areas did not respond to sleep deprivation in the same way. For example, cortical neurons favored genes associated with kinesin binding (and thus anterograde synaptic vesicle transport towards dendrites), whereas hypothalamic neurons responded by altering syntaxin-1 and calcium-dependent protein binding, consistent with their role in vesicle exocytosis (Figure S3B). In contrast, brainstem neurons displayed alterations in genes associated with cytochrome c oxidase activity, suggesting that mitochondrial energy balance is a key process altered in this area. These data suggest distinct functions of sleep in each brain region.

Interestingly, even other “support” cells in the brain exhibited region-specific functional changes. Endothelial cells showed enrichment of protein folding, and epithelial cell proliferation and migration terms, whereas microglia showed changes in genes linked to neuroinflammatory responses and glial cell activation (Figure S4C-D). We also found that EPCs, pericytes and VSMCAs exhibited significant alterations in annotations associated with protein folding, cellular transition, metal ion homeostasis, and amine metabolism (Figure S4E-G). Some of these annotations (e.g., protein folding and neuroinflammatory response) have been reported in previous sleep studies mapping bulk gene expression changes in the brain (Cirelli et al., 2004; Mackiewicz et al., 2007; Wadhwa et al., 2017). Overall, these results show that alteration of sleep need causes profound and functionally distinct gene expression changes within individual cells residing in different brain regions, which was not known before.

### Validation of transcriptional changes occurring during sleep treatments

Next, we independently validated our sequencing results by using RNAscope *in situ* hybridization assays (Figure 5A). We analyzed mRNA counts per cell in the cortex and hypothalamus of mice brain subjected to sleep treatments. Both sequencing and *in situ* hybridization results reveal that sleep deprivation enhances expression of *Mt1, Tsc22d3*, and *Gjb6* in the cortex, and expression levels decrease after recovery sleep (Figure 5B-D). Similarly, in the hypothalamus, sleep deprivation increases the expression of *Mt1*, but moderately decreases *Atp1b2*, and these levels are further decreased following recovery sleep (Figure 5E-F). The similar gene expression patterns across sleep treatments revealed by single cell sequencing and RNAscope thus validates the sequencing results (Figure 5).

**Figure 5.**
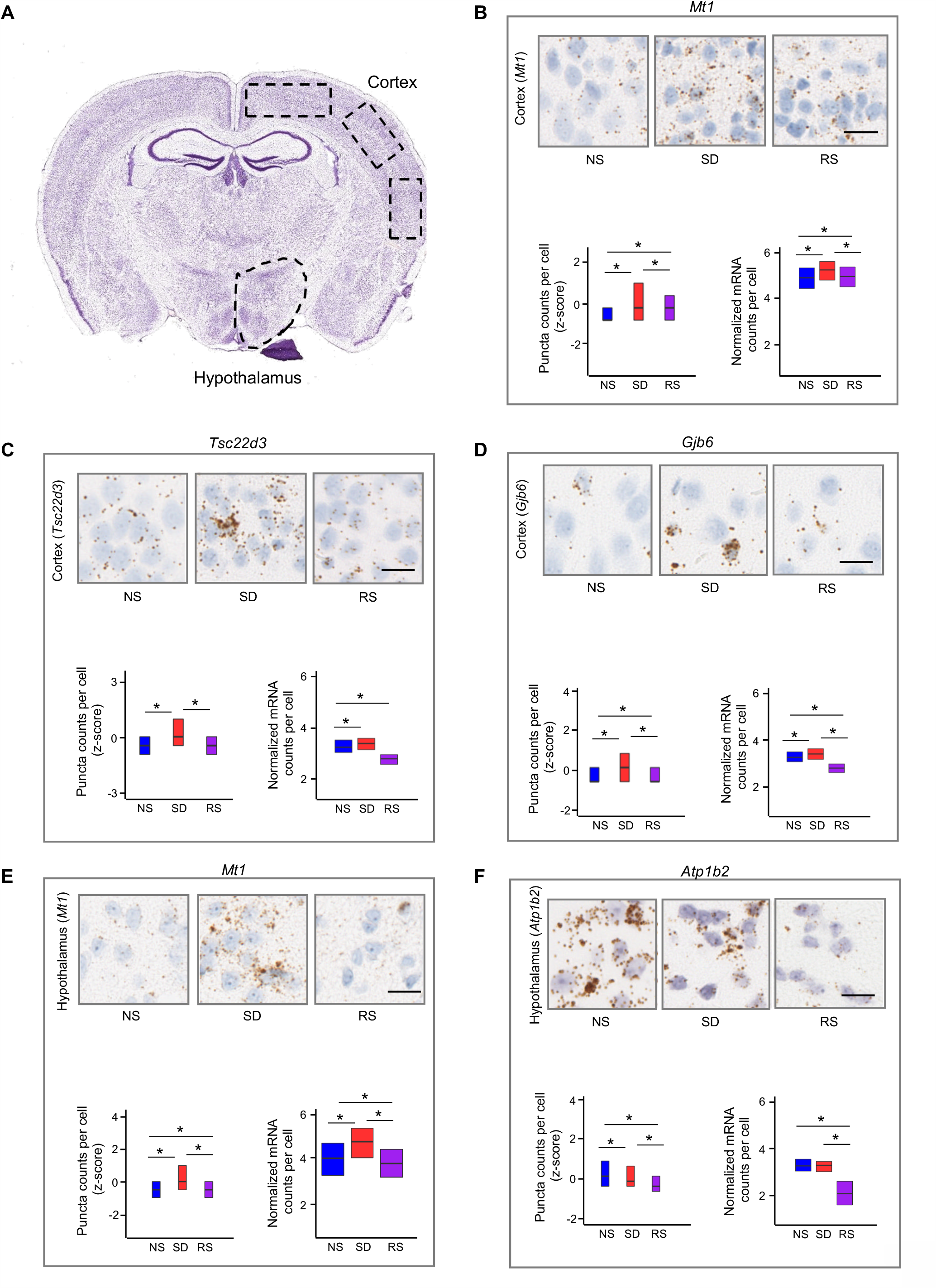
*In situ* hybridisation validation of transcriptional changes following sleep treatments. (A) Regions of interests (ROIs) for RNAscope analysis shown on coronal micrograph (from Allen Brain Atlas). (B-F) Comparison of RNAscope single cell *in situ* hybridization and sequencing of (B) *Mt1*, (C) *Tsc22d3*, (D) *Gjb6* expression in cortex, and (E) *Mt1*, (F) *Atp1b2* expression in hypothalamus across sleep treatment groups. Micrographs of mouse cortices and hypothalamic shown. mRNA molecules are brown dots as revealed by Diaminobenzidine (DAB), and nuclei are counterstained blue (with haematoxylin). Scale bar, 20µm. mRNA were counted from cells of n=3 biological replicate brains per group. Kruskal–Wallis test followed by post-hoc Dunn test. * *P* < 0.01. NS: Normal Sleep, SD: Sleep Deprivation, RS: Recovery Sleep. For sample sizes and statistics, Table S1.

### Sleep need alters astrocyte-neuron interactions

Astrocyte-neuron signaling is a key player in the maintenance of homeostatic brain function because of coupling between these cell types (Mederos et al., 2018). This is likely to be highly relevant in the context of sleep homeostasis (Halassa et al., 2010; Petit and Magistretti, 2015). We therefore used our single cell transcriptomics data to investigate sleep-related changes in astrocyte-neuron communication. We built comprehensive interactive networks of ligands from astrocytes, and receptors expressed in neurons, and *vice versa*, across all sleep treatment groups in brainstem, cortex, and hypothalamus (Figures 6A-F and S5A-G). We found significant network interactions for Apoe (Figures 6A-F) and Pomc signaling Figures 6D-E), which are key communication nodes in the hypothalamus. *Apoe* was also upregulated in the brainstem following sleep deprivation (Figure S5A). Interestingly, these genes are linked to sleep consolidation, cognition, and Alzheimer’s disease (Aibar et al., 2017; Lim et al., 2013). We also found that BDNF signaling is a mediator of crosstalk between neurons and astrocytes in the cortex (Figures S5F-G). Again, the role of *Bdnf* in sleep, insomnia and stress-related disorders has been documented (Schmitt et al., 2016). Collectively, these results suggest that specific routes of intercellular crosstalk between astrocytes and neurons are important in sleep regulation.

**Figure 6.**
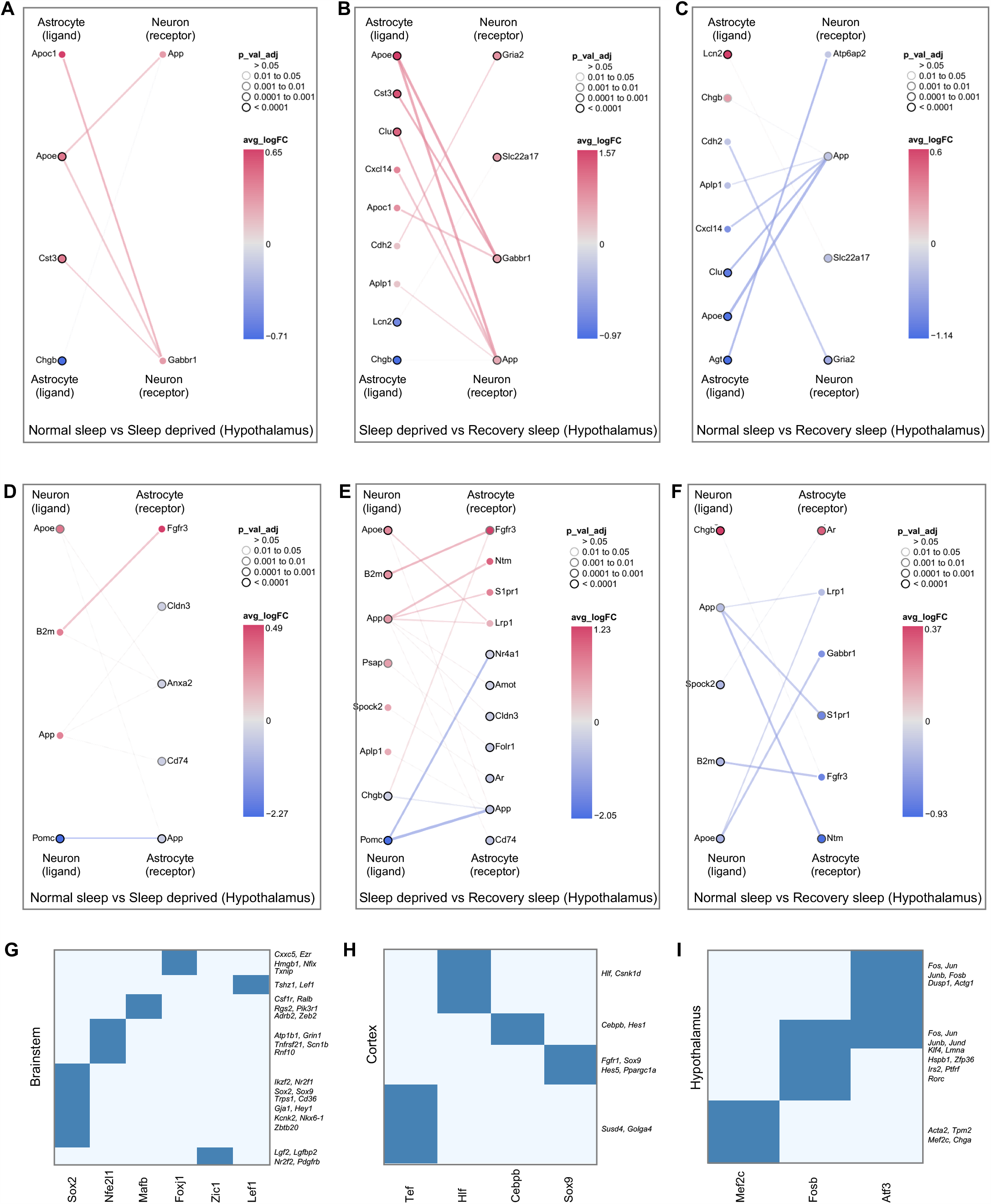
Sleep need modulates cell-cell communication and gene regulatory networks. (A-F) Interactions between differentially expressed ligands of astrocytes and receptors of neurons for (A) NS versus SD (B) SD versus RS and (C) NS versus RS comparisons, and ligands of neurons and receptors of astrocytes for (D) NS versus SD, (E) SD versus RS and (F) NS versus RS comparisons in hypothalamus. Nodes represent ligands or receptors in astrocytes and neurons, respectively. Node outline thickness indicates level of significance (Benjamini-Hochberg adjusted *P*-value). Colour of nodes and edges represent magnitude of alteration in expression (log fold-change, logFC) (see STAR Methods). (G-I) Uniquely expressed transcription factors (rows) in sleep deprived cells and their co-expressed gene sets (columns) in (G) brainstem, (H) cortex and (I) hypothalamus. Gene sets for a specific GO term shown on the right in italics. See also Figure S5 and Table S5.

### Sleep modulates gene regulatory processes

To examine how sleep state controls gene expression changes, we used SCENIC to determine regulons that might coordinate expression patterns (Aibar et al., 2017). We first evaluated regulon activity scores to profile cells under normal sleep and sleep deprived conditions. We then used regulon target information to sort transcription factors (TFs) with their sets of co-expressed target genes (Figures 6G-I). Importantly, out of all the TFs regulating gene expression in cells from sleep-deprived animals, we did not find any TF that was common to any two brain regions. Furthermore, when we performed gene-annotation enrichment analysis of co-expressed target genes (Table S5), we determined different functional annotations predominated in each brain region. The brainstem showed negative regulation of macromolecule biosynthesis (mediated by Sox2) and RNA metabolic processes (Foxj1) (Figure 6G). In contrast, cortex expresses TFs that regulate biological rhythms (Hlf) and haemopoiesis (Cebpb) (Figure 6H). Interestingly, these functions have been reported in recent sleep studies (Curie et al., 2015; McAlpine et al., 2019). Unlike other brain areas, we identified hypothalamic TFs that regulate calcium ion homeostasis (Atf3), cell death (Fosb), and muscle system process (Mef2c) (Figure 6I). Recent evidence suggests that Mef2c plays a role sleep regulation (Bjorness et al., 2020). Together, these results show that distinct TFs in each brain region appear to regulate single cell gene expression.

### Sleep need regulates translational responses of cortical astrocytes and neurons

Multiple studies in several biological domains have shown that there is a poor relationship between gene and protein expression in global analyses (Abreu et al., 2009; Maier et al., 2009; Mauvoisin et al., 2014; Reddy et al., 2006; Robles et al., 2014; Schwanhäusser et al., 2011). Moreover, in the context of sleep, no studies have been performed to investigate sleep need regulation within specific cell types in the cortex (or any other brain region). Given that cortex has two principal cell types that are likely to be key in sleep regulation (astrocytes and neurons), we next examined protein expression in these cells. We purified astrocytes and neurons from cortex using magnetic cell separation (Figures 1A, 7A, see STAR Methods) and then performed quantitative global- and phospho-proteomics by using multiplex tandem mass tag (TMT) labelling coupled with liquid chromatography–tandem mass spectrometry (LC–MS/MS) (Figure 7A, see STAR Methods).

**Figure 7.**
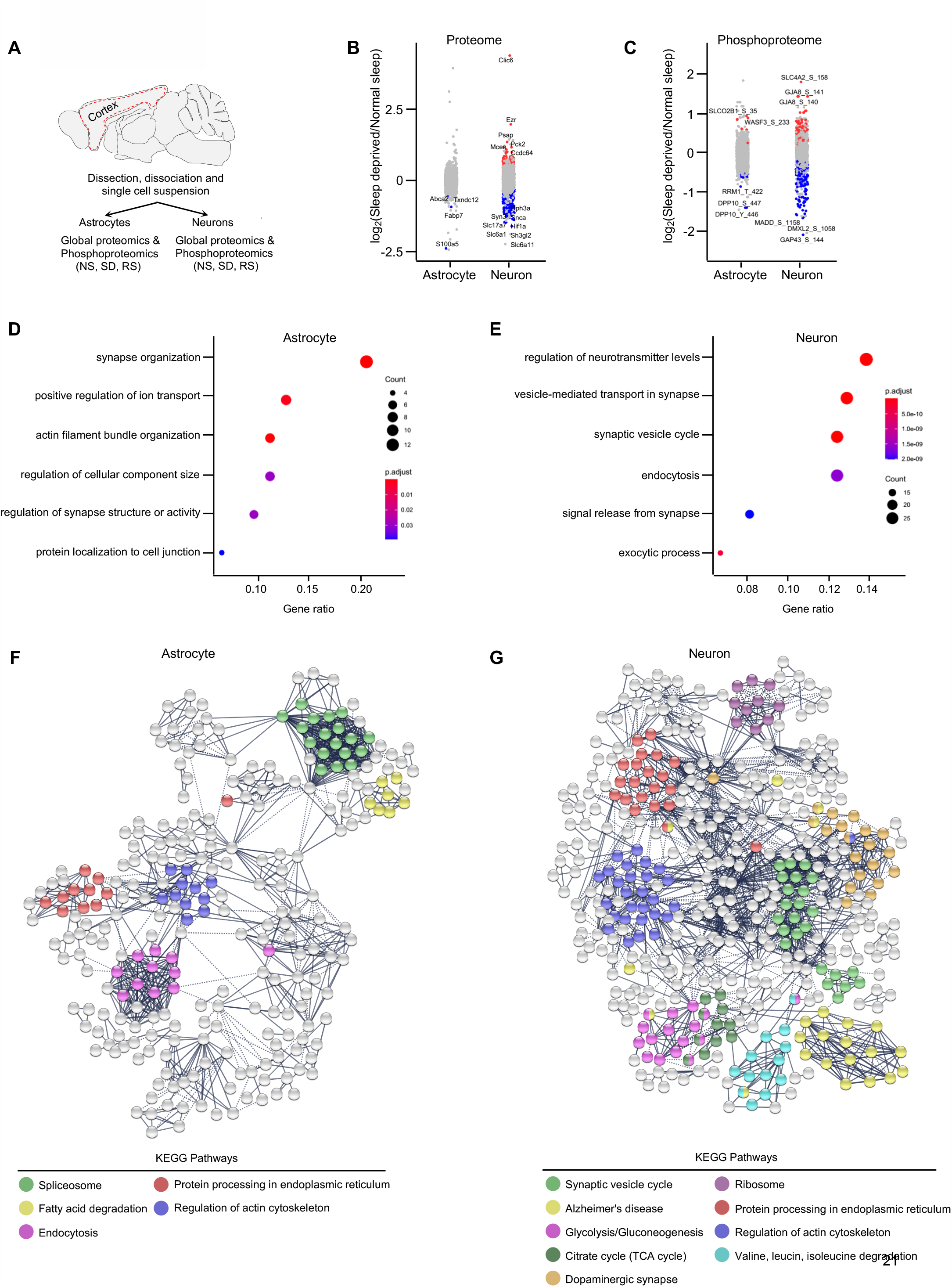
Sleep deprivation regulates the global and phosphoproteome of astrocytes and neurons in cerebral cortex. (A) Schematic of proteomics workflow. Dotted red line indicates area of cortex dissected for magnetic cell separation. (B and C) Strip charts showing changes in (B) global and (C) phosphoproteome for Sleep Deprivation (SD)/Normal Sleep (NS) comparisons in astrocytes and neurons. Unpaired *t*-tests followed by false discovery rate (FDR) analysis were used to compare groups. Significantly upregulated and downregulated protein expression is color coded with red and blue, respectively (Benjamini-Hochberg adjusted *P*-value < 0.1). Proteins in grey are not significantly changed after sleep deprivation. (D and E) Dot plots showing significantly enriched top functional annotations from global proteomes of (D) astrocytes and (E) neurons. Circle size is proportional to protein count and level of significance (Benjamini-Hochberg adjusted *P*-value < 0.05) is colour coded. (F and G) Interaction networks of significantly altered (FDR < 0.1) global proteins in (F) astrocytes and (G) neurons across sleep treatments. Functional enrichment of proteins (KEGG pathways) in the network is colour coded. The networks were generated using STRING Version 11.0 database (PPI enrichment *P*-value < 0.001). See also Figures S6 and S7, and Tables S6, S7, S8, and S9.

We quantified 2,454 proteins expressed in astrocytes and 1,401 in neurons. To understand how sleep-wake states affect protein abundance in astrocytes and neurons, we compared protein expression in both cell types exposed to different sleep treatments (Figures 7B, S6A and S6C). The proportion of altered proteins was higher in neurons (318/1,401, 22.7%; 1.5-fold cutoff, FDR < 0.1) than in astrocytes (93/2,454, 3.7%; 1.5-fold cutoff, FDR < 0.1) following sleep treatments. Interestingly, we saw similar transcriptional changes when we compared cortical astrocytes and neurons (Table S3). We hypothesized that this similarity might be a manifestation of astrocyte-neuron communication to maintain synaptic and metabolic homeostasis, as astrocytes bolster the structure and function of neurons. To test this, we performed gene ontology (GO) analysis on differentially expressed proteins in both cell types. In astrocytes, we found prominent alterations of actin filament organization, regulation of cellular component size, synapse organization and regulation of synapse structure or activity. In contrast, vesicle-mediated transport in synapse, regulation of neurotransmitter levels, and signal release from synapse were enriched in neurons (Figures 7D and 7E). Next, we determined these differences are due to sleep treatments, not due to a global difference in protein abundance between the cell types, for this we analyzed the gene annotations of all proteins expressed versus differentially expressed proteins within these cell types (Tables S6 and S7).

We also analyzed protein-protein interaction (PPI) networks using STRING to identify biological pathways altered by sleep treatments in astrocytes and neurons (Szklarczyk et al., 2019) (Figures 7F-G). Sleep-regulated astrocyte proteins segregated into clusters enriched for the spliceosome, fatty acid degradation and endocytosis (Figure 7F). Neurons, however, were focused on various networks including the synaptic vesicle cycle, glucose metabolism (glycolysis, gluconeogenesis, TCA cycle), and branched chain amino acid (valine, leucine, isoleucine) degradation (Figure 7G). Thus, neurons display prominent responses of vesicular machinery and energy production, in contrast to support function responses of astrocytes. Interestingly, proteins associated with the endoplasmic reticulum and regulation of actin cytoskeleton function were enriched in both cell types, suggesting a concerted role in sleep.

### Sleep need regulates post-translational responses of cortical astrocytes and neurons

Protein phosphorylation modulates protein function in reversible manner (Humphrey et al., 2015). Recent evidence suggests that sleep need modulates protein phosphorylation cycles(Brüning et al., 2019; Wang et al., 2018), but cell type contributions to these are unknown. We therefore performed phosphoproteomics on neuron and astrocyte lysates (Figure 7A, see STAR Methods) and compared protein phosphorylation profiles of sleep treatment groups in both cell types (Figures 7C, S6B and S6D). We identified 1591 phosphorylation sites in astrocytes and 1159 in neurons. Sleep treatments altered 134 phosphorylation sites in astrocytes and 263 in neurons (Figures 7C, S6B and S6D). These phosphorylation sites were from 92 and 156 unique proteins from astrocytes and neurons, respectively.

Next, we determined the overlap between sleep related changes in protein phosphorylation from whole brain proteome data from Wang et al. (2018) and our cortical astrocyte and neuron data and were able to assign cell types to some of the differentially expressed phosphosites found previously (Wang et al., 2018) (Figure 7C, Table S10). Comparative GO analysis between total versus differentially altered phosphorylated proteins revealed that sleep treatments alter phosphorylation of proteins associated with the regulation of cellular component size, protein complex assembly and neurotransmitter secretion in astrocytes. By contrast, phosphorylation affects the synaptic vesicle cycle, vesicle-mediated transport in synapses, and calcium mediated signaling proteins in neurons (Tables S8-9). Thus, like single cell transcriptomics, global proteomics and phosphoproteomics demonstrate that sleep need regulates cellular functions in astrocytes and neurons in a cell-specific manner.

## Discussion

In this study, we provide the first comprehensive single cell transcriptome profiling of the cell types within three brain areas (brainstem, cortex, and hypothalamus) of mice in different sleep stages. We found sleep need modulates transcriptional patterning of astrocytes, neurons, endothelial cells, and microglia in all three brain regions. We also utilized our single cell gene expression information to interrogate remodeling of astrocyte-neuron signaling during sleep and determined expression changes in transcription factors that may mediate these effects. Cell-specific proteomics reveals that sleep need regulates translational and post-translational responses of astrocytes and neurons in the cortex differently.

Previous bulk RNA quantification methods could not reveal the cell-specific differential expression changes that we found in this study. Our differential gene expression analysis within individual cell types did not show a universal sleep-related changes, but rather a distinct pattern of transcriptional responses of sleep need in each cell type. Moreover, this was not the same for different brain regions, further highlighting that cells in their local milieu respond to the same stimulus (sleep deprivation and recovery) in different ways. This had not been appreciated before because bulk methods by their nature average out the individual contributions of diverse cell types in each brain region. Indeed, in our analyses, we were able to retrospectively assign gene expression changes seen in previous bulk microarray or sequencing studies to specific cell types, enhancing the utility of these studies further.

Single cell RNA-seq analysis has also advanced our understanding of the functional pathways affected by sleep perturbations. Pathway analysis revealed different pattern of sleep driven changes across cell types and brain areas. For example, we found that sleep need alters gene expression associated with ribosomal biogenesis in astrocytes, but vesicle mediated transport in synapses and nucleotide/nucleoside phosphate metabolism in neurons. Endothelial cells exhibit changes in protein folding pathways, whereas microglia show neuroinflammatory responses. Interestingly, protein folding and neuroinflammatory responses have been documented in previous sleep studies (Cirelli et al., 2004; Korin et al., 2019; Mackiewicz et al., 2007; Wadhwa et al., 2017). Our results provide cell specificity to these responses in the brain for the first time. Furthermore, the same cell type in different brain areas performed variable molecular functions to regulate diverse biological processes, indicating brain region specific molecular responses of sleep need. Collectively, these data indicate that the transcriptional response of sleep loss is not identical in all cell types and brain areas, rather this response is cell type, and brain region specific.

How are cell specific transcriptional changes brought about? We were able to identify transcription factors (TF) that mediate gene expression changes in each cell type in sleep deprived cells from the brainstem, cortex, and hypothalamus. Interestingly, we found no common TF in any two brain areas. This analysis revealed that the key TFs in brainstem are Sox2, Nfe2l1, Mafb, Foxj1, Zic1, and Lef1, whereas in cortex they were Tef1, Hlf1, Cebpb, and Sox9, and in hypothalamus Mef2c, Fosb and Atf3. Future studies will explore directly (e.g., using loss-of-function approaches) the roles of these TFs in sleep homeostasis.

Astrocytes and neurons are major cells types of the brain that are likely to be important in sleep. Since we were able to delineate transcriptional changes in each of these cell types for the first time with single cell RNA-seq, this enabled us to determine novel relationships between the two cell types in sleep. We generated networks of receptor-ligand interactions to decipher sleep driven astrocyte-neuron communications in all three brain regions. The significance of these interactive networks is of high importance as some recent findings show that astrocytes interact with neurons to modulate sleep circuitry, and metabolic coupling between them, across the sleep-wake cycle (Fellin et al., 2012; Halassa et al., 2010; Petit and Magistretti, 2015). Thus, the discovery of novel factors, and their source and target, is a fruitful avenue to define novel therapeutic interventions of sleep abnormalities. We found Apoe, Pomc and Bdnf signaling were modulated by sleep in the different brain regions. Of note, the role of these genes in sleep related pathology have been described (Goldstein et al., 2018; Lim et al., 2013; Schmitt et al., 2016). Our study advances our understanding by showing the cell types involved, and the directionality of changes in intercellular ligand-receptor signaling. Such insights will enable further mechanistic studies targeting these ligands and receptors to modulate sleep, and potentially develop novel therapeutic agents that manipulate these pathways.

To gain further insights into the functional implication of sleep need in astrocytes and neurons, we performed proteomics and phosphoproteomics. We found that 22% of neuronal proteins responded to sleep perturbation compared to only 3% in astrocytes. Interestingly, a previous sleep study using a 6 h sleep deprivation paradigm concluded that the whole brain proteome remains globally stable following sleep treatments (Wang et al., 2018). There are some possible explanations of the disparity between those findings and ours. First, differences could be due to reciprocal changes in different brain areas and/or cell types that may have been “masked” by bulk cell proteomics, particularly in whole brain samples; our study used cortex only. Second, 6 h of sleep deprivation (compared to 12 h used here) may not have been sufficient time for transcriptional changes to become apparent at the protein level. We propose that astrocytes show relative resistance to sleep need and may thereby provide a stable support to rapidly changing neurons via metabolic coupling (Petit and Magistretti, 2015). The biological functions of sleep-mediated alterations include actin filament bundle organization, positive regulation of ion transport, synapse organization, and fatty acid degradation in astrocytes, and vesicle-mediated transport in synapse, regulation of neurotransmitter levels, and glucose metabolism in neurons. These functions support the proposed importance of synapse and energy homoeostasis as functions of sleep-wake cycles (Benington and Heller, 1995; Tononi and Cirelli, 2014). Our results now provide cell-specific compartmentalization of these functions.

We found that changes in cell-specific gene and protein (and phosphoprotein) expression profiles were not correlated. This indicates that sleep is regulated differentially at transcriptional, translational, and post-translational levels within individual cell types. The ensemble of cellular processes regulated at these levels include synapse organization, synaptic vesicle recycling, protein folding, ribosome function, and glucose metabolism. As such, we propose that prolonged wakefulness during sleep deprivation leads not only to sleepiness, but could also impair cognition, physiology, and metabolism via the diverse molecular responses we identified in individual brain cells. Moreover, we suggest that sleep need could be a compensatory mechanism to reverse such molecular changes in the brain. Overall, by using scRNA-seq and cell-specific proteomics we provide a resource detailing the molecular responses of sleep need at the individual cell level. These data will help to advance a variety of efforts towards understanding the cellular and molecular mechanisms of sleep and wake and provide a critical path to dissect sleep functions.

## Acknowledgments

A.B.R. acknowledges funding from the Perelman School of Medicine, University of Pennsylvania and the Institute for Translational Medicine and Therapeutics (ITMAT) at the University of Pennsylvania.

## Author contributions

A.B.R., P.K.J. and U.K.V. conceived and designed the experiments. P.K.J. and M.N. performed sleep phenotyping by EEG/EMG analysis. P.K.J. performed mouse sleep deprivation experiments and tissue dissections. P.K.J. and U.K.V. performed single cell isolation and library preparation. U.K.V., P.K.J. and A.B.R. analyzed single cell transcriptomics data. P.K.J and U.K.V performed RNAScope *in situ* hybridization analysis. P.K.J. and U.K.V. performed astrocyte and neuron isolation from mouse cortex. S.R and U.K.V. performed quantitative proteomics and phosphoproteomics. S.R., P.K.J., U.K.V., and A.B.R. analyzed proteomics data. A.B.R. supervised the entire study and secured funding. The manuscript was written by P.K.J. and A.B.R, with input from U.K.V. All authors agreed on the interpretation of data and approved the final version of the manuscript.

## MATERIALS & METHODS

### Experimental model and subject details

All animal studies were carried out in concordance with an approved protocol from Institutional Animal Care and Use Committee (IACUC) at Perelman School of Medicine at the University of Pennsylvania, or under license by the United Kingdom Home Office under the Animals (Scientific Procedures) Act 1986, with Local Ethical Review by the Francis Crick Institute Animal Welfare & Ethical Review Body Standing Committee (AWERB). Wild type, male C57BL/6J mice were purchased from Charles River and allowed to acclimatize in the animal unit for at least 2 weeks prior to experiments. Mice were selected such that, at the time of experiments, they were aged between 9-11 weeks. Before sleep deprivation, mice were singly housed in automated sleep fragmentation chambers (Campden/Lafayette Instruments Model 80391) for habituation with *ad libitum* access to food and water under standard humidity and temperature (21±1°C) on a 12-h light: 12-h dark cycle (Kaushal et al., 2012a).

### Sleep deprivation

Sleep deprivation were performed in the sleep fragmentation home cages as previously demonstrated and adapted to new model 80391 having the possibility of much faster sweep time (Kaushal et al., 2012b). Briefly, this device act by applying tactile stimulus with horizontal bar sweeping just above the cage floor (bedding) from one side to the other side of cage. Once sweeper is on, animals need to step over approaching sweeper to resume their unrestrained behaviors. Sleep deprivation was initiated at lights on [Zeitgeber Time 0 (ZT0)] by switching on the motors, choosing the continuous sweeping mode (approximately 7.5 seconds cycle time) and stopped after 12 hours at lights off (ZT12). To assess how 12-h of sleep deprivation affected the sleep/wake behaviors, 3 days baseline EEG/EMG were recorded after mice were acclimatized for a week. Mice were recorded from for next 3-4 days (which encompassed sleep deprivation and recovery phases). Post sleep deprivation, animals were allowed to recover for 24-h. Baseline recording 1-day before sleep deprivation was used as the control condition (normal sleep). Following sacrifice by cervical dislocation, whole brains were isolated at ZT12 (i.e. lights off) for *ad libitum* Normal Sleep (NS, ZT0-12), Sleep Deprivation (SD, ZT0-12) and 12-h sleep deprivation followed by 24-h of Recovery Sleep (RS, ZT12-12) groups. Isolated whole brains were placed in ice-cold Hibernate EB (BrainBits LLC, HEB) media and brainstem, cortex and hypothalamus were quickly dissected, and tissue collected for preparation of single cell suspensions.

### Sleep phenotyping

Mice aged between 9-11 weeks were implanted with a telemetry transmitter (HD-X02, Data Sciences International, St. Paul, MN, USA) connected to electrodes for continuous EEG (Electroencephalography)/EMG (Electromyography) recording. Under anaesthesia (isoflurane; induction 3-4%, maintenance 2-2.5%), two stainless steel EEG electrodes (length of screw shaft: 2.4 mm; head diameter: 2.16 mm; shaft diameter: 1.19 mm; Plastics One, Roanoke, VA, USA) were implanted epidurally over the right frontal and parietal cortices, as previously described (Hasan et al., 2011). The electrodes were connected to the telemetry transmitter via medical-grade stainless steel wires. The EEG electrodes were covered with dental cement (Kemdent, Purton, Swindon, UK). Two EMG stainless-steel leads were inserted into the neck muscles ∼5 mm apart and sutured into place. The telemetry transmitter was placed in a subcutaneous pocket and positioned along the left dorsal flank. Analgesia was administered at the onset of the surgery (subcutaneous injection of buprenorphine (Vetergesic) at 0.1 mg/kg and meloxicam (Metacam) at 10 mg/kg). Animals were allowed to recover for a minimum of 10 days before being subjected to experimental protocols. EEG/EMG signals were recorded continuously for 6-7 days using Data Sciences International hardware and Dataquest ART software (Data Sciences International, St. Paul, MN, USA). The EEG/EMG data were transmitted at 455 kHz to an RPC-1 receiver (Data Science International) and sampled at 250 Hz.

### Single cell suspensions and library preparation

The protocol for obtaining single cell suspensions for all three brain regions was adapted from that used by Holt and colleagues (Holt and Olsen, 2016). Briefly, tissue dissections were dissociated with Papain (BrainBits LLC, PAP/HE) following the manufacturer’s instruction followed by manual trituration using fire polished Silanized Pasteur pipette and filtration through 70µm cell strainer (Miltenyi Biotec, 130-095-823). Cells were pelleted at 300 × g, 10 minutes, the supernatant carefully removed, and cells resuspended in a minimal volume of PBS with 0.5% of BSA (Sigma-Aldrich, A7906). To reduce debris, we incubated the cell suspensions with myelin removal beads (Miltenyi Biotec, 130-096-733) and passed this through a LS Column (Miltenyi Biotec, 130-042-401) with 70µm pre-separation filter. Flow-through contained unlabelled cells, which were collected and counted (Countess II Automated Cell Counter; Cat # AMQAX1000) estimate yield and viability. Library preparation was carried out with a 10X Genomics Chromium Single Cell Kit Version 2. Suspensions were prepared as described above and diluted in PBS with 0.5% of BSA to concentration ∼ 350 cells/ µl and added to 10X Chromium RT mix to achieve a loading target between 7,000-12,000 cells. After cell capture, downstream cDNA synthesis, library preparation, and sequencing were performed according to manufacturer’s instructions (10X Genomics).

### Astrocyte and neuron separation from cortical single cell suspension

To perform cell type specific proteomics and phosphoproteomics, astrocytes and neurons were separated from cortical single cell suspensions as previously described (Holt and Olsen, 2016) with slight modifications. Single cell suspensions were incubated with FcR Blocking Reagent (Miltenyi Biotec, 130-092-575) and Anti-ACSA-2 MicroBeads (Miltenyi Biotec, 130-097-678) for 10-15 minutes at 2-8°C. Cells were spun down for 10 minutes at 300 × g at room temperature and resuspended in a minimal volume of PBS with 0.5% of BSA before passing through a LS Column. Flow-through was collected for further neuron separation, and the LS Column that retained astrocytes was removed from the magnetic field and the ACSA-2 labelled astrocytes were eluted. Cells were pelleted from flow-through at 300 × g, 8 minutes, the supernatant carefully removed, and resuspended in a minimal volume of PBS with 0.5% of BSA. Suspensions were incubated with Non-Neuronal Cell Biotin-Ab cocktail (Miltenyi Biotec, 130-115-389) for 5 minutes and Anti-Biotin MicroBeads (Miltenyi Biotec, 130-090-485) for 10 minutes at 2-8°C, respectively. Following incubation, suspensions were passed through a LS Column placed in a magnetic field and non-labelled neuronal cells were collected in flow-through.

### TMT-based quantitative proteomics and phosphoproteomics

Cell lysis and protein extraction were performed in the presence of protease and phosphatase inhibitors. Protein precipitation was performed with 1:6 volume of pre-chilled (-20°C) acetone for overnight at 4°C. After overnight incubation, lysates were centrifuged at 14,000 g for 15 minutes at 4°C. Supernatants were discarded without disturbing pellets, and pellets were air-dried for 2-3 minutes to remove residual acetone. Then pellets were dissolved in 300 µL 100mM TEAB buffer. Sample processing for TMT-based quantitative proteomics and phosphoproteomics was performed following the same protocol as described previously (Ray et al., 2019). In brief, protein concentration in each sample was determined by using the Pierce™ BCA Protein Assay Kit (Thermo Fisher Scientific, 23225). 75µg protein per condition was transferred into new microcentrifuge tubes and 5 µL of the 200 mM TCEP was added to reduce the cysteine residues and the samples were then incubated at 55°C for 1-h. Subsequently, the reduced proteins were alkylated with 375 mM iodoacetamide (freshly prepared in 100 mM TEAB) for 30 minutes in the dark at room temperature. Then, trypsin (Trypsin Gold, Mass Spectrometry Grade; Promega, V5280) was added at a 1:40 (trypsin: protein) ratio and samples were incubated at 37°C for 12-h for proteolytic digestion. After in-solution digestion, peptide samples were labelled with 10-plex TMT Isobaric Label Reagents (Thermo Fisher Scientific, 90113) following the manufacturer’s instructions. The reactions were quenched using 5µL of 5% hydroxylamine for 30 minutes. Proteins from astrocytes or neurons for the three experimental conditions, i.e. NS, SD, and RS (3 biological replicates for each) were labeled with the nine TMT reagents within a TMT 10-plex reagent set, while the 10th reagent was used for labeling an internal pool containing an equal amount of proteins from each sample. Application of multiplexed TMT reagents allowed comparison of NS, SD, and RS samples within the same MS run, which eliminated the possibility of run-to-run (or batch) variations.

In order to perform phosphoproteome analysis, TMT labelled peptides were subjected to TiO2-based phosphopeptide enrichment according to the manufacturer’s instructions (High-Select™ TiO2 Phosphopeptide Enrichment Kit, Thermo Fisher Scientific, A32993). The flow-through and washes from the TiO2 enrichment step were combined, evaporated to dryness and subjected to Fe-NTA-based phosphopeptide enrichment according to the manufacturer’s instructions (High-Select™ Fe-NTA Phosphopeptide Enrichment Kit, Thermo Fisher Scientific, A32992). Phosphopeptides enriched using TiO2- and Fe-NTA-based methods were combined, evaporated to dryness. TMT-labelled samples (both for global and phosphoproteomics analysis) was resuspended in 5% formic acid and then desalted using a SepPak cartridge according to the manufacturer’s instructions (Waters, Milford, Massachusetts, USA). Eluate from the SepPak cartridge was evaporated to dryness and resuspended in buffer A (20 mM ammonium hydroxide, pH 10) prior to fractionation by high pH reversed-phase (RP) chromatography using an Ultimate 3000 liquid chromatography system (Thermo Scientific). In brief, the sample was loaded onto an XBridge BEH C18 Column (130Å, 3.5 µm, 2.1 mm X 150 mm, Waters, UK) in buffer A and peptides eluted with an increasing gradient of buffer B (20 mM Ammonium Hydroxide in acetonitrile, pH 10) from 0-95% over 60 minutes. The resulting fractions (8 fractions per sample) were evaporated to dryness and resuspended in 1% formic acid prior to analysis by nano-LC MSMS using an Orbitrap Fusion Lumos mass spectrometer (Thermo Scientific).

### Nano-LC Mass Spectrometry

High pH RP fractions were further fractionated using an Ultimate 3000 nano-LC system in line with an Orbitrap Fusion Lumos mass spectrometer (Thermo Scientific). In brief, peptides in 1% (vol/vol) formic acid were injected onto an Acclaim PepMap C18 nano-trap column (Thermo Scientific). After washing with 0.5% (vol/vol) acetonitrile 0.1% (vol/vol) formic acid peptides were resolved on a 250 mm × 75 μm Acclaim PepMap C18 reverse phase analytical column (Thermo Scientific) over a 150 minutes organic gradient, using 7 gradient segments (1-6% solvent B over 1 minutes, 6-15% B over 58 minutes, 15-32%B over 58 minutes, 32-40%B over 5 minutes, 40-90%B over 1 minutes, held at 90%B for 6 minutes and then reduced to 1%B over 1 minutes) with a flow rate of 300nL min^−1^. Solvent A was 0.1% formic acid and Solvent B was aqueous 80% acetonitrile in 0.1% formic acid. Peptides were ionized by nano-electrospray ionization at 2.0kV using a stainless-steel emitter with an internal diameter of 30 μm (Thermo Scientific) and a capillary temperature of 275°C. All spectra were acquired using an Orbitrap Fusion Tribrid mass spectrometer controlled by Xcalibur 4.1 software (Thermo Scientific) and operated in data-dependent acquisition mode using an SPS-MS3 workflow. FTMS1 spectra were collected at a resolution of 120,000, with an automatic gain control (AGC) target of 200,000 and a max injection time of 50 ms. Precursors were filtered with an intensity threshold of 5000, according to charge state (to include charge states 2-7) and with monoisotopic peak determination set to Peptide. Previously interrogated precursors were excluded using a dynamic window (60s +/-10ppm). The MS2 precursors were isolated with a quadrupole isolation window of 0.7m/z. ITMS2 spectra were collected with an AGC target of 10 000, max injection time of 70 ms and CID collision energy of 35%. For FTMS3 analysis, the Orbitrap was operated at 50,000 resolution with an AGC target of 50,000 and a max injection time of 105 ms. Precursors were fragmented by high energy collision dissociation (HCD) at a normalised collision energy of 60% to ensure maximal TMT reporter ion yield. Synchronous Precursor Selection (SPS) was enabled to include up to 5 MS2 fragment ions in the FTMS3 scan.

### RNAscope in situ hybridization

Brains were quickly isolated and fixed in 10% neutral-buffered formalin (NBF) for 24-h, followed by ethanol gradient dehydration and infiltration with melted paraffin in an automated processor. An array of 4 µm coronal sections of whole brain was prepared, in order to collect different regions of cortex, and RNAscope *in situ* hybridization was performed according to manufacturer’s reference guide, using the single-plex RNAscope® assay kit-BROWN (Advanced Cell Diagnostics) on a Leica Biosystems BOND RX platform, as described previously (Anderson et al., 2016). Brain slices were baked and deparaffinized on the instrument, followed by target retrieval and protease treatment. Hybridization was performed by using probe *Mt1* (ACD, 547711), *Tsc22d3* (ACD, 448341), *Gjb6* (ACD, 458811) and *Atp1b2* (ACD, 417131) followed by amplification, DAB chromogenic detection and counterstain with haematoxylin. Imaging of sections was carried out on a Leica Biosystems Aperio AT2 Digital Pathology Scanner and analysis performed using Visiopharm software.

### Sleep scoring and spectral analysis

Vigilance states for consecutive 4 sec epochs were classified by visual inspection of the EEG and EMG signals, according to standard criteria (Franken et al., 1998), as follows: wakefulness (high and variable EMG activity and a low amplitude EEG signal), non-rapid eye movement sleep (NREMS; high EEG amplitude, dominated by slow waves and low amplitude EMG), and rapid eye movement sleep (REMS; low EEG amplitude, theta oscillations of 5-9 Hz, and loss of EMG muscle tone). EEG power spectra were computed for consecutive 4 sec epochs using Welch’s method in Matlab 2017a (MathWorks, Natick, MA, USA; frequency range, 0.25-25 Hz; resolution, 0.25 Hz; Hanning window function). Epochs containing EEG artefacts were discarded from EEG spectral analyses (% of recording time: 7.9 ± 0.8%). EEG power spectra were determined for NREMS, REMS, and wakefulness during the 12-h light-dark periods of the 3 recording days. EEG power spectra are expressed as a percentage of total EEG power (frequency range: 1-25 Hz; resolution: 1 Hz). EEG delta power total and during NREMS were computed by adding the EEG power in the frequencies ranging from 1 to 4.5 Hz. Averaged over consecutive intervals to which an equal number of 4-second NREM sleep epochs contribute, and then expressed as a percentage of levels reached between ZT8-12 during the day 1 (baseline) (Franken et al., 2001).

### Analysis of single cell RNA-seq data

FASTQ files containing sequence reads were mapped to the mouse reference genome GRCm38 using Cell Ranger Version 2.2.0 on a Linux high-performance computing system. The output of the Cell Ranger pipeline (filtered counts) was parsed into R version 4.0.0 to perform downstream analysis with the various R packages – Seurat version 3.1.5 (Butler et al., 2018), Monocle version 2.10.1 (Trapnell et al., 2014) and SCENIC version 1.1.2.2 (Aibar et al., 2017). The data were first normalized and log-transformed using Seurat function NormalizeData. Differential gene expression was computed by Wilcoxon rank sum test method available in the Seurat function FindMarkers. Cluster analysis on the first 13 principal components was performed after calculating the JackStraw. Clusters were visualized with uniform manifold approximation and projection (UMAP) in Seurat with default settings. Seurat functions UMAPPlot, VlnPlot, FeaturePlot, DotPlot, and DoHeatmap were adapted for visualization of gene expression. All Monocle and SCENIC analyses were performed on a Linux computing system using 256 GB RAM spread across 32 compute cores. The expression matrix from the Seurat data object was then used to perform Single Cell regulatory Network Inference and Clustering analysis with SCENIC. The species-specific RcisTarget database of motif rankings was downloaded from https://resources.aertslab.org/cistarget/databases/mus_musculus/mm10/refseq_r80/mc9nr/. The standard SCENIC workflow was run on each dataset individually or on a merged data together as described at http://scenic.aertslab.org.

### Determination of cell identity clusters

To determine the cell-type identity of each cluster from the brainstem, cortex and hypothalamus, we used multiple cell-type specific/enriched markers genes that have been previously described in the single cell transcriptomics of mouse brain in the literature (Chen et al., 2017; Mickelsen et al., 2019; Saunders et al., 2018; Zeisel et al., 2018). Each cluster showing high expression level of a known markers specific to a particular cell were considered as cluster of that cell-type (Table S2). In general, we selected the threshold of average fold change above 2-3 and false discovery rate (FDR) < 0.001. We also cross validated the other clusters from same brain area for the absence of these markers. We found the available markers were sufficient to define all major cell types in all three brain regions, with one “unknown” (unclassifiable) cell-type in brainstem.

### Astrocyte-neuron communication networks

Astrocyte-neuron communication networks were predicted by previously described intercellular communication networks (Kirouac et al., 2010; Ximerakis et al., 2019). The cell interactomes for differentially expressed genes were created based on ligand-receptor interactions. In the networks, nodes represent ligands and receptors expressed in astrocytes and neurons and, vice-versa. The border of nodes represents level of significance (FDR). Edges represent interactions between them, and the color of nodes and edges represents the magnitude of differential expression.

### Database search and statistical analysis of quantitative proteomics data

Quantitative proteomics and phosphoproteomics raw data files were analysed using the MaxQuant computational platform (*version* 1.5.2.8) with the Andromeda search engine (Cox and Mann, 2008). MS2/MS3 spectra were searched against UniProt database specifying *Mus musculus* (Mouse) taxonomy (Proteome ID: UP000000589; Organism ID: 10090; Protein count: 52026). All searches were performed using “Reporter ion MS3” with “10-plex TMT” as isobaric labels with a static modification for cysteine alkylation (carbamidomethylation), and oxidation of methionine (M) and protein N-terminal acetylation as the variable modifications. Phospho (STY) was included as an additional variable modification for the phosphopeptide enrichment analyses. Trypsin digestion with maximum two missed cleavages, minimum peptide length as seven amino acids, precursor ion mass tolerance of 5 ppm and fragment mass tolerance of 0.02 Da were specified in all analyses. The false discovery rate (FDR) was specified at 0.01 or 0.05 for peptide spectrum match (PSM), protein and site decoy fraction. TMT signals were corrected for isotope impurities based on the manufacturer’s instructions. Subsequent processing and statistical analysis of quantitative proteomics and phosphoproteomics datasets were performed using Perseus (*version* 1.5.5.3) (Tyanova et al., 2016). During data processing, reverse and contaminant database hits and candidates identified only by site were filtered out. For differential quantitative proteomics analyses, categorical annotation (NS/SD/RS) was applied to group reporter ion intensities, values were Log2 transformed, and were normalized by “subtract mean (column-wise)” in each TMT reporter ion channel. Proteins groups were filtered for valid values (at least 80% in each group). Unpaired t-tests followed by false discovery rate (FDR) analysis was performed to compare sleep treatments groups.

### Gene ontology (GO) analysis

Significantly and differentially expressed transcripts and proteins across sleep treatment groups were subjected to GO analysis. We performed overrepresentation analysis (Boyle et al., 2004) that were implemented in clusterProfiler (Yu, 2018). Significantly enriched GO terms were visualized as dot plots and enrichment maps.

### Network Analysis

To investigate association between significant and differentially expressed global proteins in sleep treatment groups from astrocytes and neurons we extracted protein associations from the Search Tool for the Retrieval of Interacting Genes or Proteins (STRING) database (Szklarczyk et al., 2019). STRING identified 462/463 significant proteins and returned 863 interactions in astrocytes (protein-protein interaction (PPI) enrichment *P*-value < 0.001), 695/701 significant proteins and 1702 interactions in neurons (PPI enrichment *P*-value < 0.001) at a confidence cut-off value 0.9 (the highest confidence level in STRING).

### Analysis of RNAscope *in situ* hybridization images

RNA markers were quantified based on dots quantified within cortical and hypothalamic areas. We counted the cells from three brains per sleep treatment group (Ximerakis et al., 2019). We counted the almost entire cortex (1500 × 800 µm, 1200 × 800 µm and 900 × 800 µm) and hypothalamus by defined region of interests (ROIs) (Figure 5A). The number of transcripts per cell was determined by using Visiopharm’s image analysis algorithm.

### Statistical methods

All experimental subjects are biological replicates. For single cell sequencing, dissections of the same anatomical region were pooled from three animals for the preparation of single cell suspensions for each group. GraphPad Prism 8 or R software was used to perform statistical tests. Kruskal– Wallis tests were performed to compare sleep treatments groups. Following two-way analysis of variance (ANOVA), Fisher’s LSD tests were performed for comparisons. Repeated measures tests were performed for same-subject comparisons. *P* < 0.05 was considered as significant. For transcriptomics and proteomics analyses, we used FDR < 0.1, unless indicated otherwise. Detailed sample sizes, statistical tests and results are reported in the figure legends and Table S1.

## Supplemental Figures

**Figure S1.**
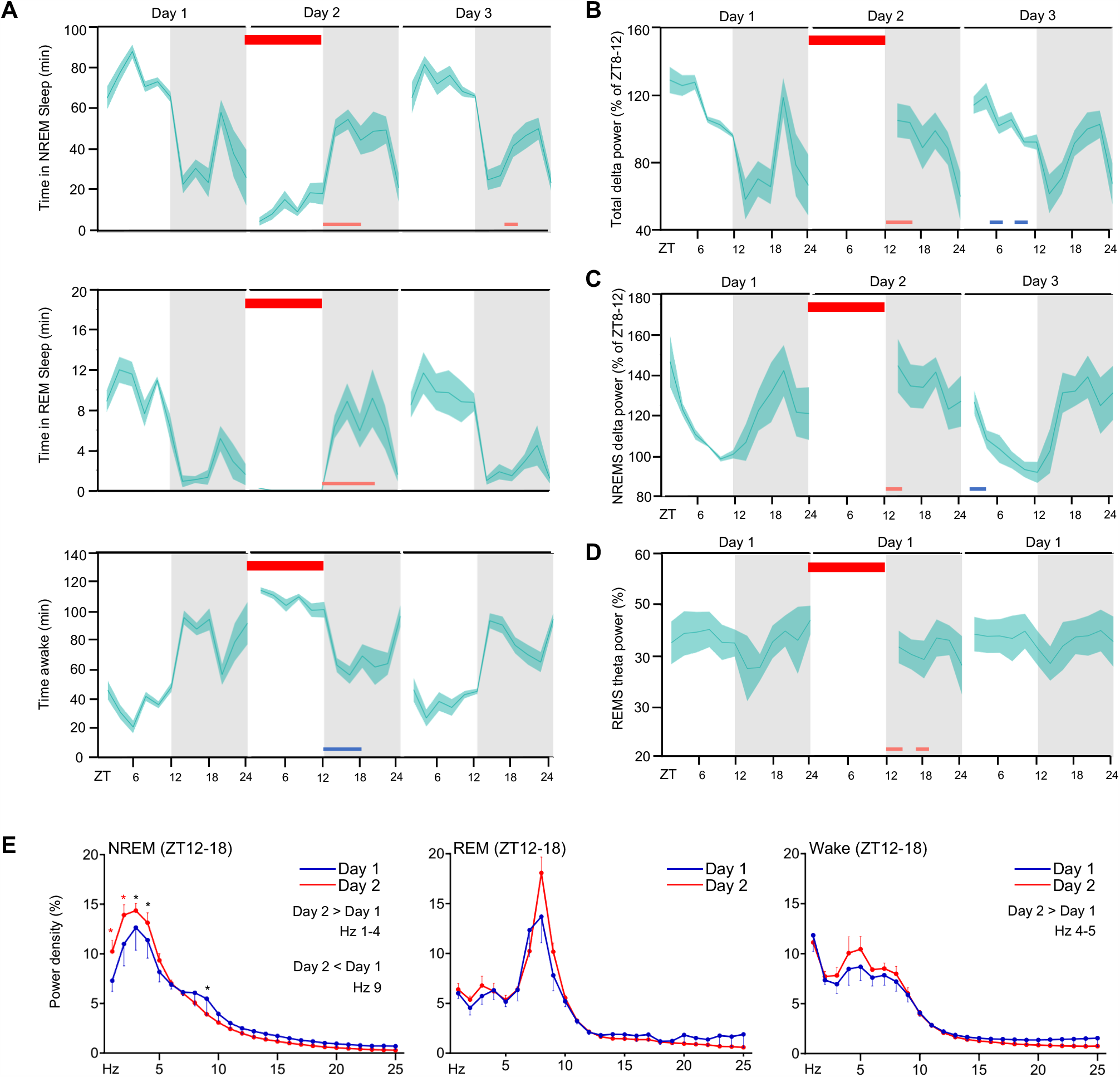
EEG validation of sleep phenotype during sleep deprivation experiment, Related to Figure 1. (A-D) Ribbon plots showing time course of (A) NREM, REM and wake states, and total (B), NREM (C) Delta (δ)-power percentage of ZT8-12 from Day 1 and REM (D) Theta (θ)-power. Horizontal red bar represents timeline of sleep deprivation (Day 2, ZT 0-12; 12 h duration). Pink and blue bars under the plots denote significant increase and decrease, respectively, compared to Day 1 (*P* < 0.05). (E) Relative EEG power spectra from ZT 12-18 on Day 1 versus Day 2. Data are mean±s.e.m.. Repeated-measures two-way ANOVA followed by post-hoc Fisher’s Least Significant Difference (LSD) test. n = 6 biological replicates, *(black), *P* < 0.01; *(red), *P* < 0.001. Sample sizes and statistics, Table S1.

**Figure S2.**
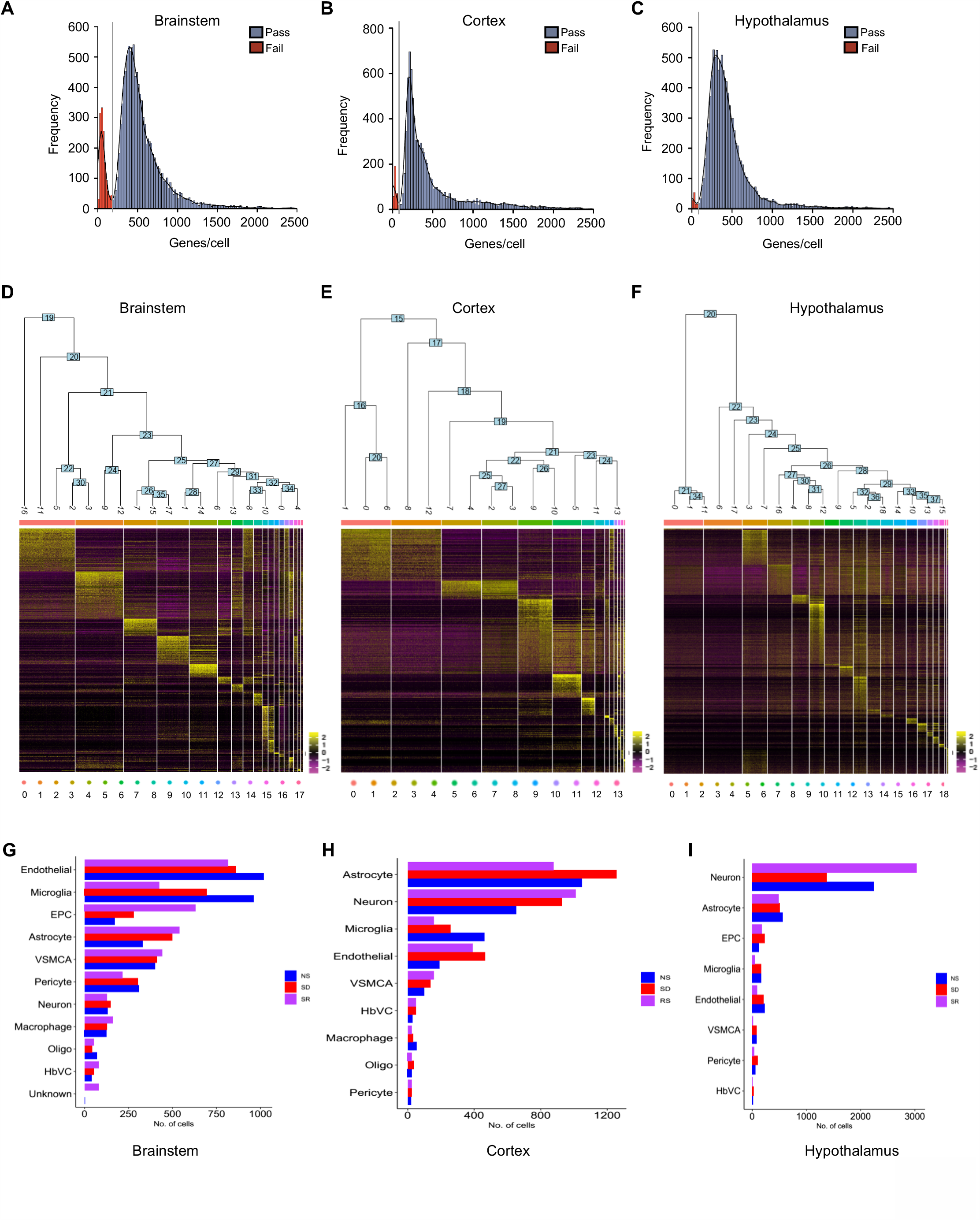
Quality control and identification of cell clusters based on specific expression patterns in brainstem, cortex and hypothalamus, Related to Figure 1, 2, 3 and 4. (A-C) Distribution of genes per cell in (A) brainstem, (B) cortex and (C) hypothalamus. (D-F) Heatmap and dendrogram showing distinct expression clusters for gene markers in (D) brainstem, (E) cortex and (F) hypothalamus. (G-I) Bar chart showing number of cell types identified across Normal sleep (NS), Sleep deprivation (SD) and Recovery Sleep (RS) in (G) brainstem, (H) cortex and (I) hypothalamus.

**Figure S3.**
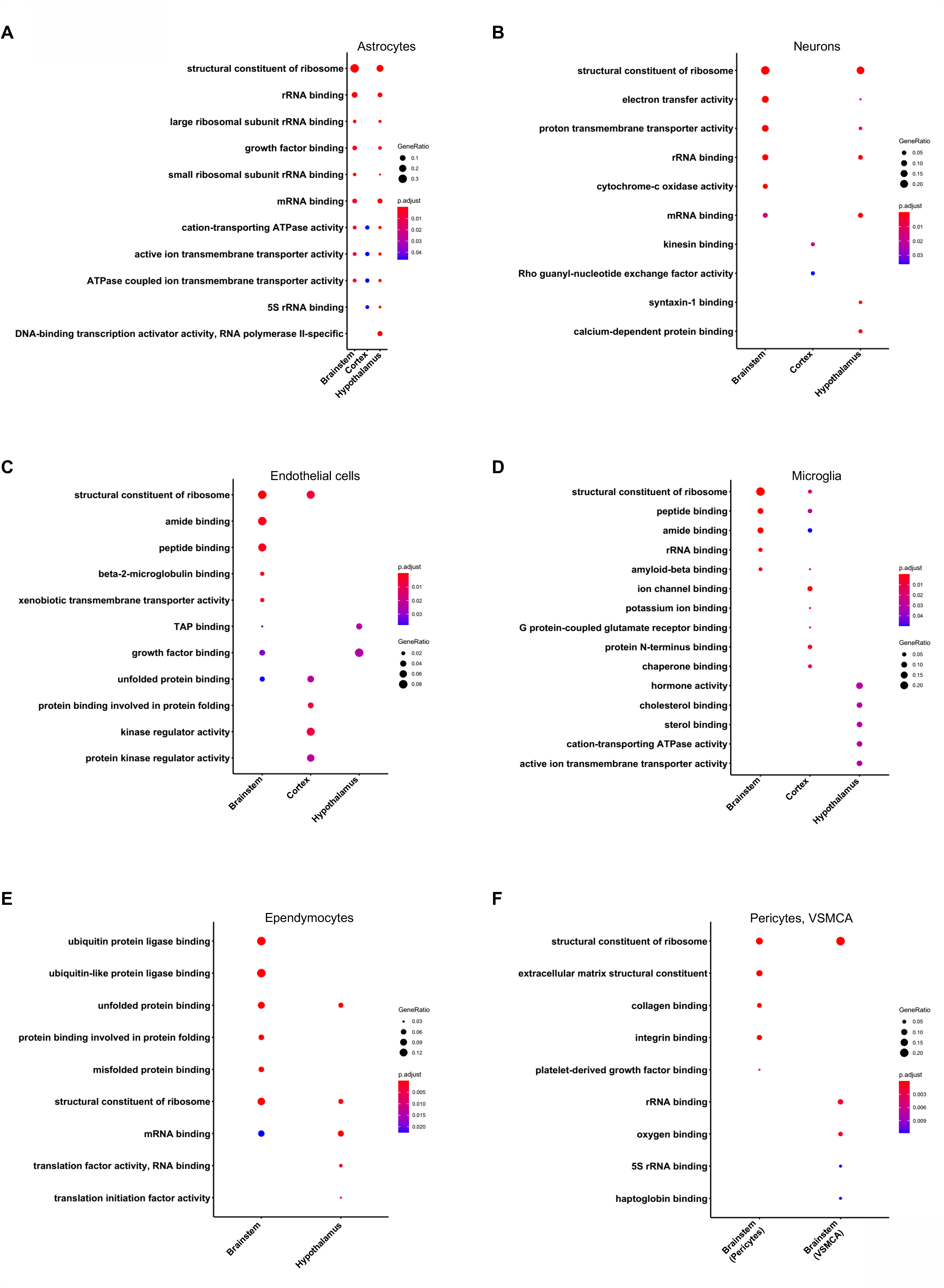
Molecular functions of differentially expressed genes in the cells from brainstem, cortex and hypothalamus across sleep treatments, Related to Figure 2, 3 and 4. (A-F) Dot plots showing comparison of significantly enriched (Benjamini-Hochberg adjusted *P*-value < 0.01) gene ontology (GO) terms of molecular functions in (A) astrocyte, (B) neuron, (C) endothelial cells, (D) microglia, (E) ependymocytes, (F) pericytes and VSMCA of all three brain regions. Circle size is proportional to gene ratio and level of significance (Benjamini-Hochberg adjusted *P*-value) is colour coded.

**Figure S4.**
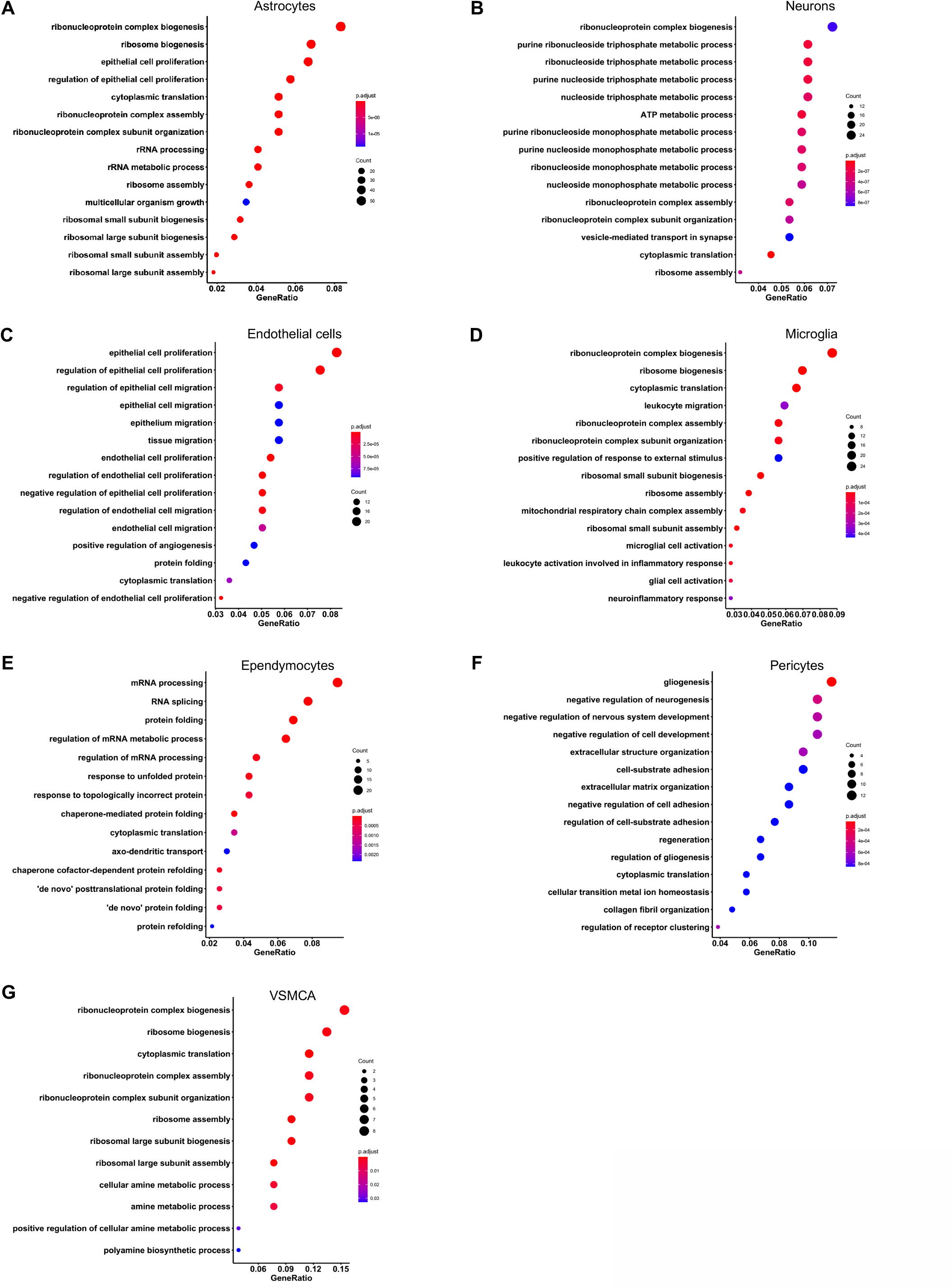
Biological functions of differentially expressed genes in the cells from brainstem, cortex and hypothalamus across sleep treatments, Related to Figure 2, 3 and 4. (A-G) Dot plots showing significantly enriched (Benjamini-Hochberg adjusted *P*-value < 0.01) gene ontology (GO) terms of biological processes in (A) astrocyte, (B) neuron, (C) endothelial cells, (D) microglia, (E) ependymocytes, (F) pericytes and (G) VSMCA. Circle size is proportional to genes count and level of significance (Benjamini-Hochberg adjusted *P*-value) is colour coded.

**Figure S5.**
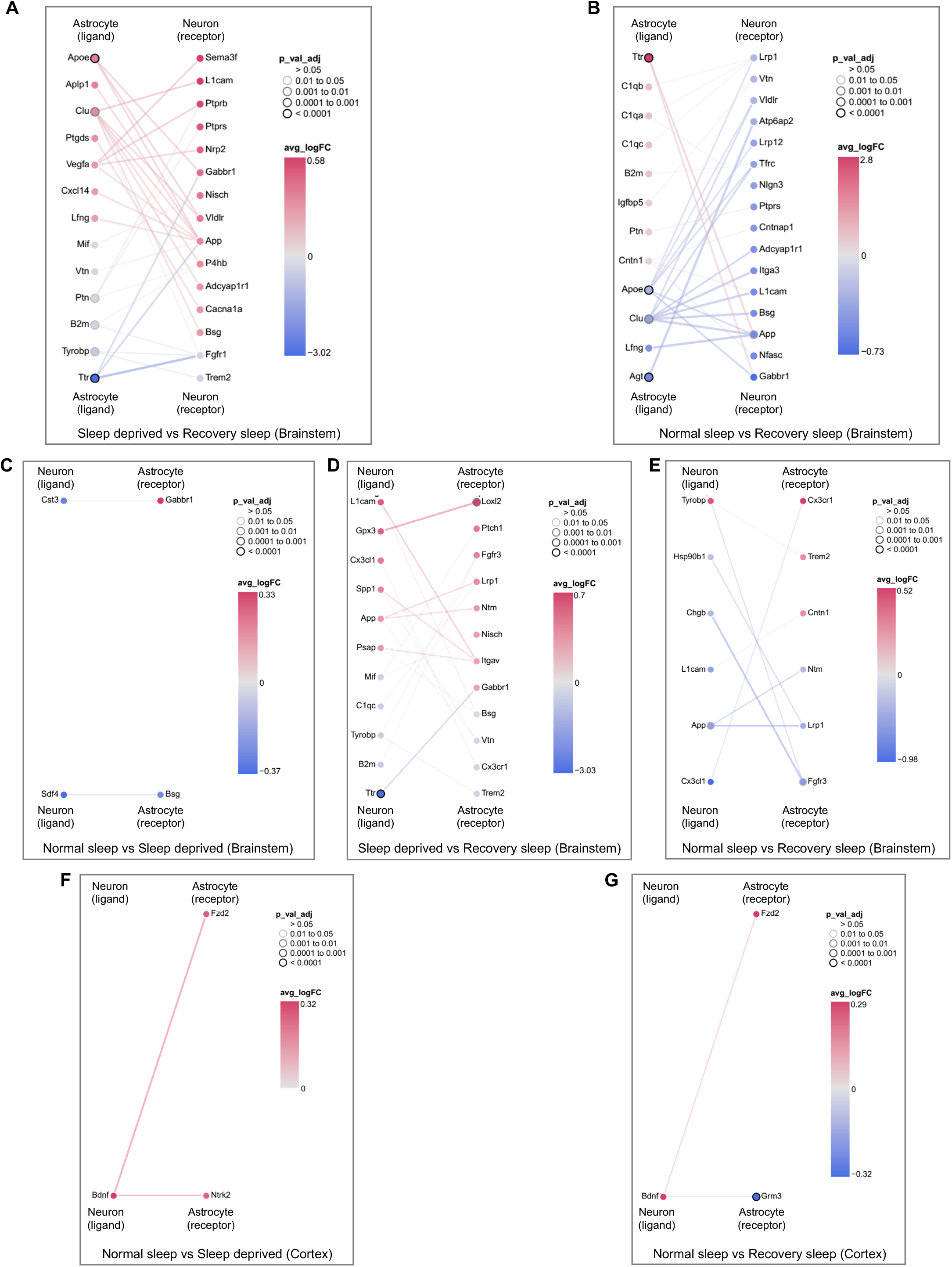
Sleep need modulates cell-cell communications networks in brainstem and cortex, related to Figure 6. (A-E) Interactions between differentially expressed ligands of astrocytes and receptors of neurons for (A) SD versus RS (B) NS versus RS and, ligands of neurons and receptors of astrocytes for (C) NS versus SD, (D) SD versus RS and (E) NS versus RS comparisons in brainstem. (F and G) Interactions between differentially expressed ligands of neurons and receptors of astrocytes for (F) NS versus SD and (G) NS versus RS comparisons in cortex. Nodes represent ligands or receptors in astrocytes and neurons, respectively. Node outline thickness indicates level of significance (Benjamini-Hochberg adjusted *P*-value). Colour of nodes and edges represent magnitude of alteration in expression (log fold-change, logFC) (see STAR Methods). NS: Normal Sleep, SD: Sleep Deprivation, RS: Recovery Sleep.

**Figure S6.**
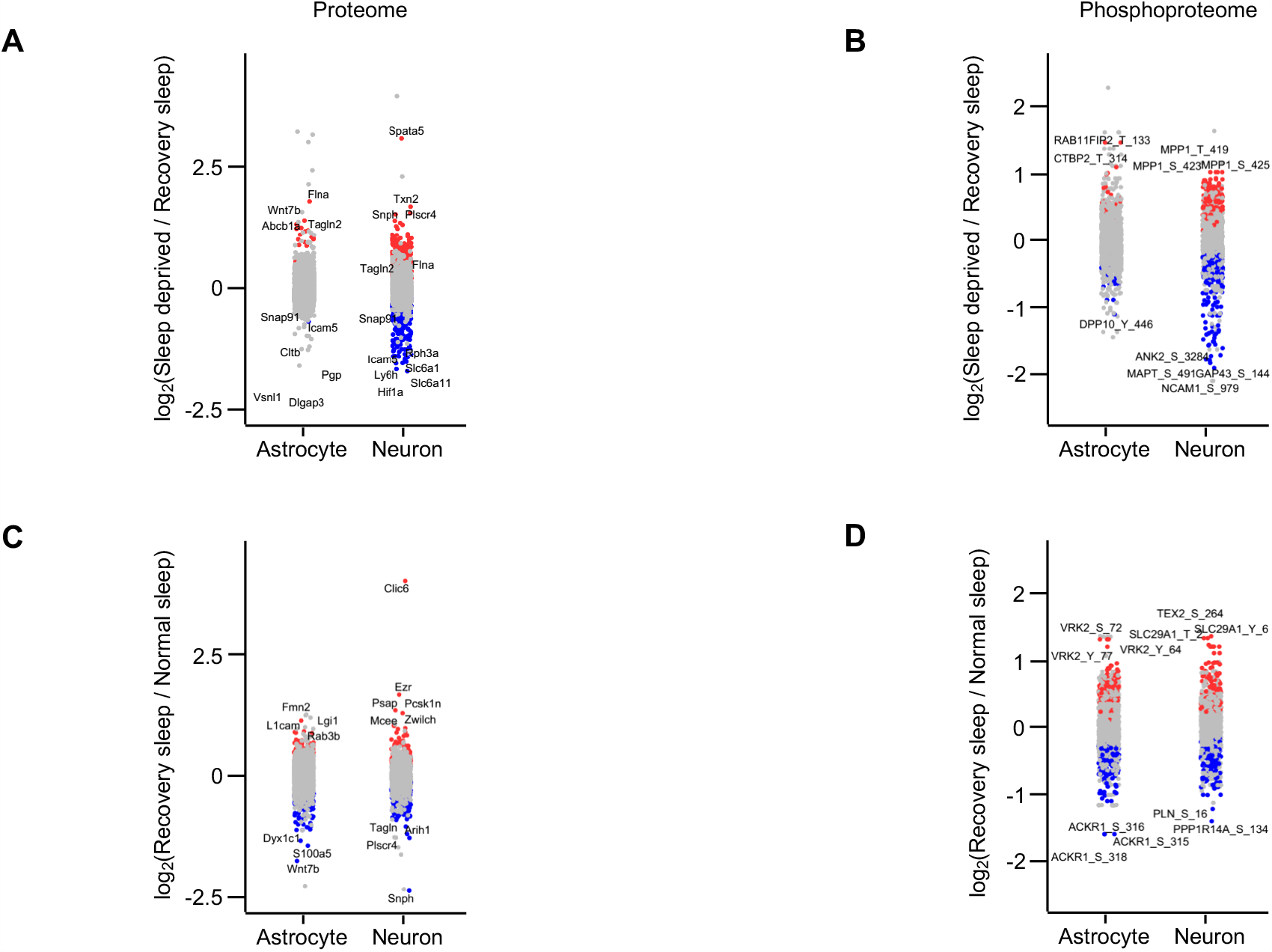
Translational and post-translational profiles of astrocytes and neurons from cortex across sleep-wake states, Related to Figure 7. (A-D) Strip chart showing changes in (A) global and (B) phosphorylated proteins for Sleep Deprivation (SD)/Recovery Sleep (RS), whereas (C) and (D) show change in global and phosphorylated proteins, respectively for Recovery Sleep (RS)/Normal Sleep (NS) comparisons. Unpaired t-tests followed by false discovery rate (FDR) analysis were used to compare groups. Significantly upregulated and downregulated protein expression is colour coded with red and blue, respectively (Benjamini-Hochberg adjusted *P*-value < 0.1). Proteins in grey are not significantly changed following sleep treatments.

**Figure S7.**
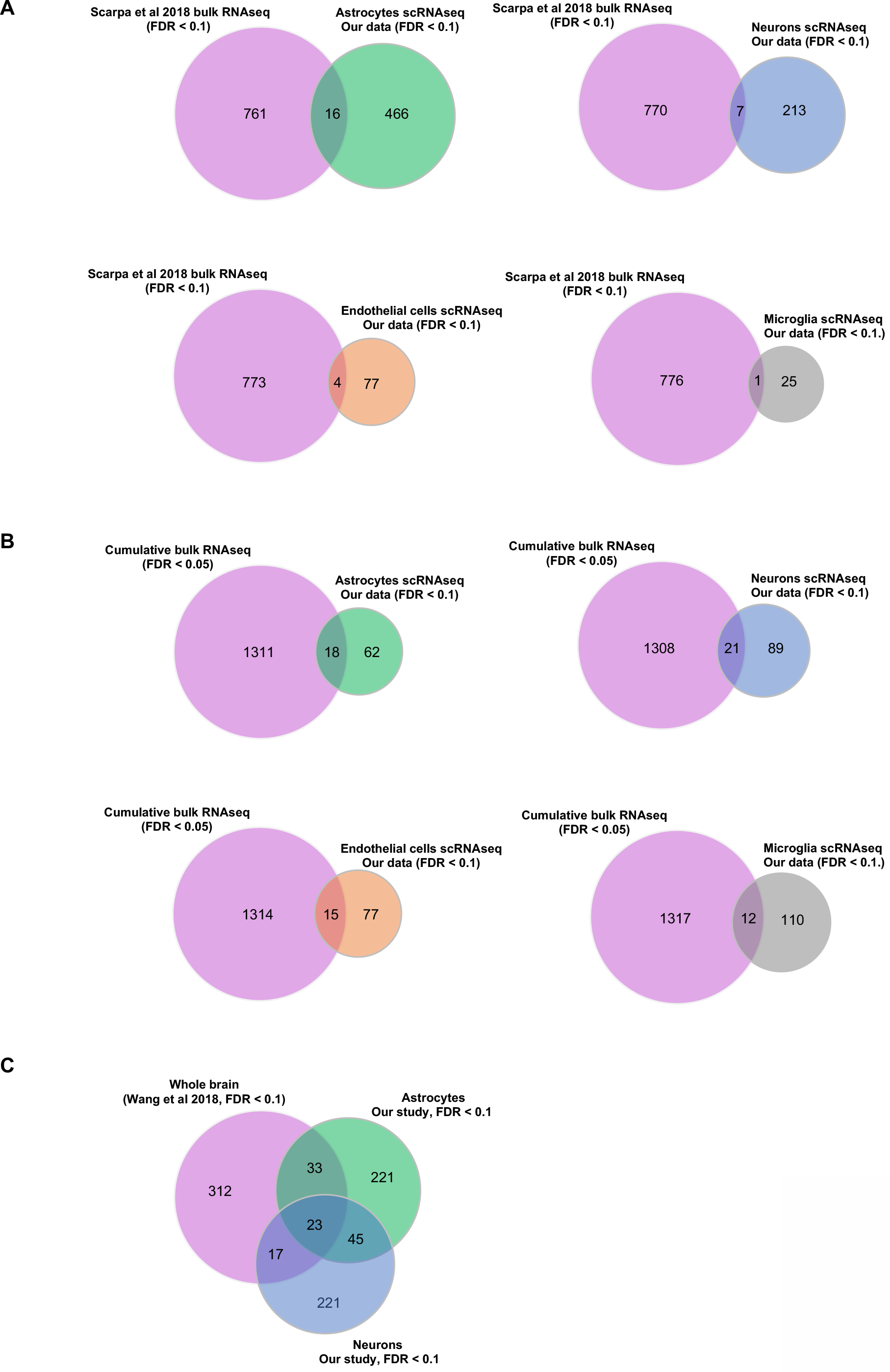
Cell-type specificity of previously published sleep related transcriptional and translational studies compared to scRNAseq results, Related to Figures 3, 4 and 7. (A and B) Venn diagrams of sleep related transcriptional alterations at cumulative bulk cell and single-cell (astrocytes, neurons, endothelial cells and microglia) in (A) cortex, and (B) hypothalamus. (C) Venn diagram of sleep driven alterations in protein phosphorylation in the whole brain and, astrocytes and neurons. Cumulative bulk cell transcription studies were Diessler et al., 2018; Gerstner et al., 2016; Scarpa et al., 2018, and whole brain protein phosphorylation data from Wang et al., 2018. See Tables S4 and S10.

## Supplemental Tables

**Table S1.**
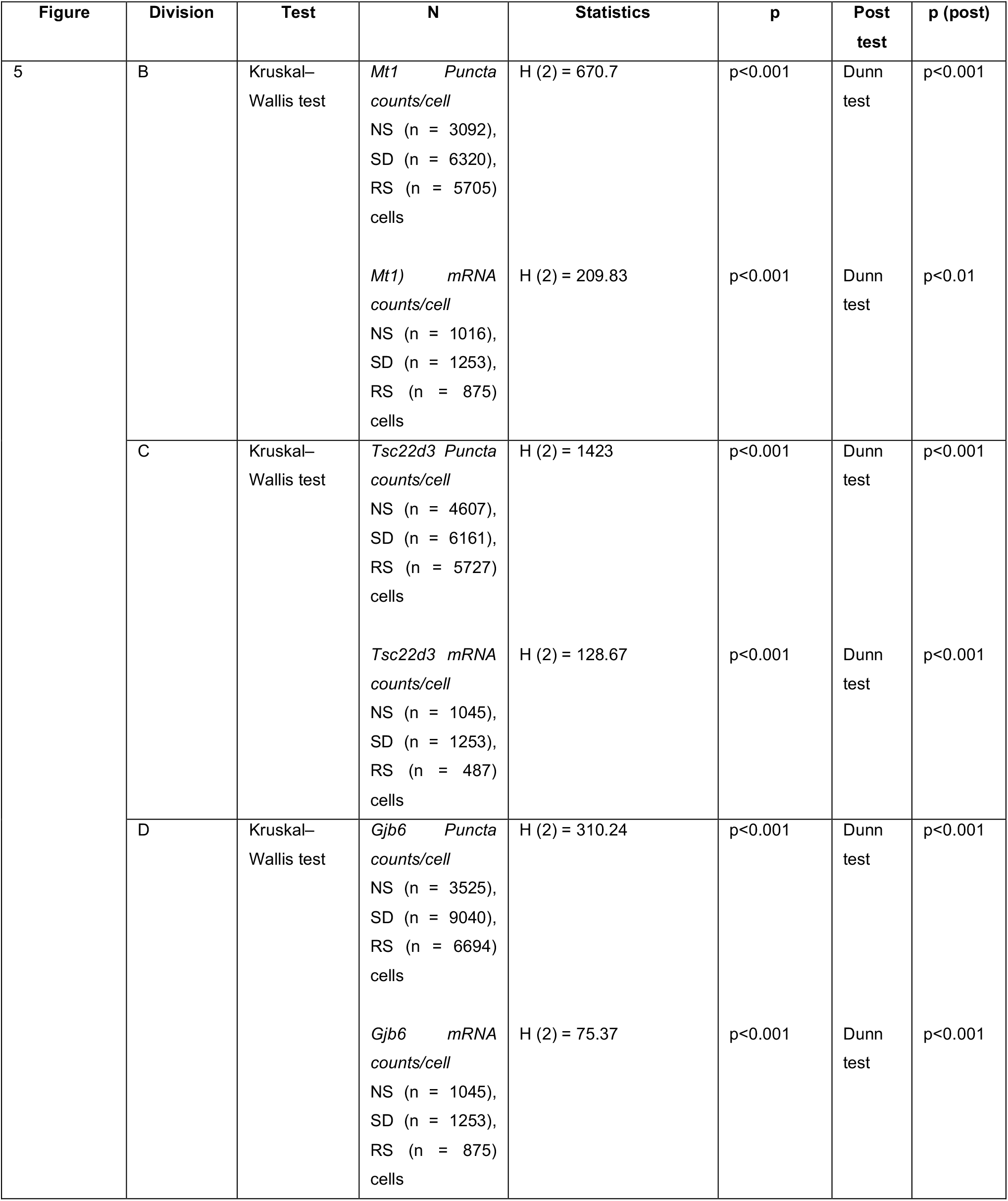
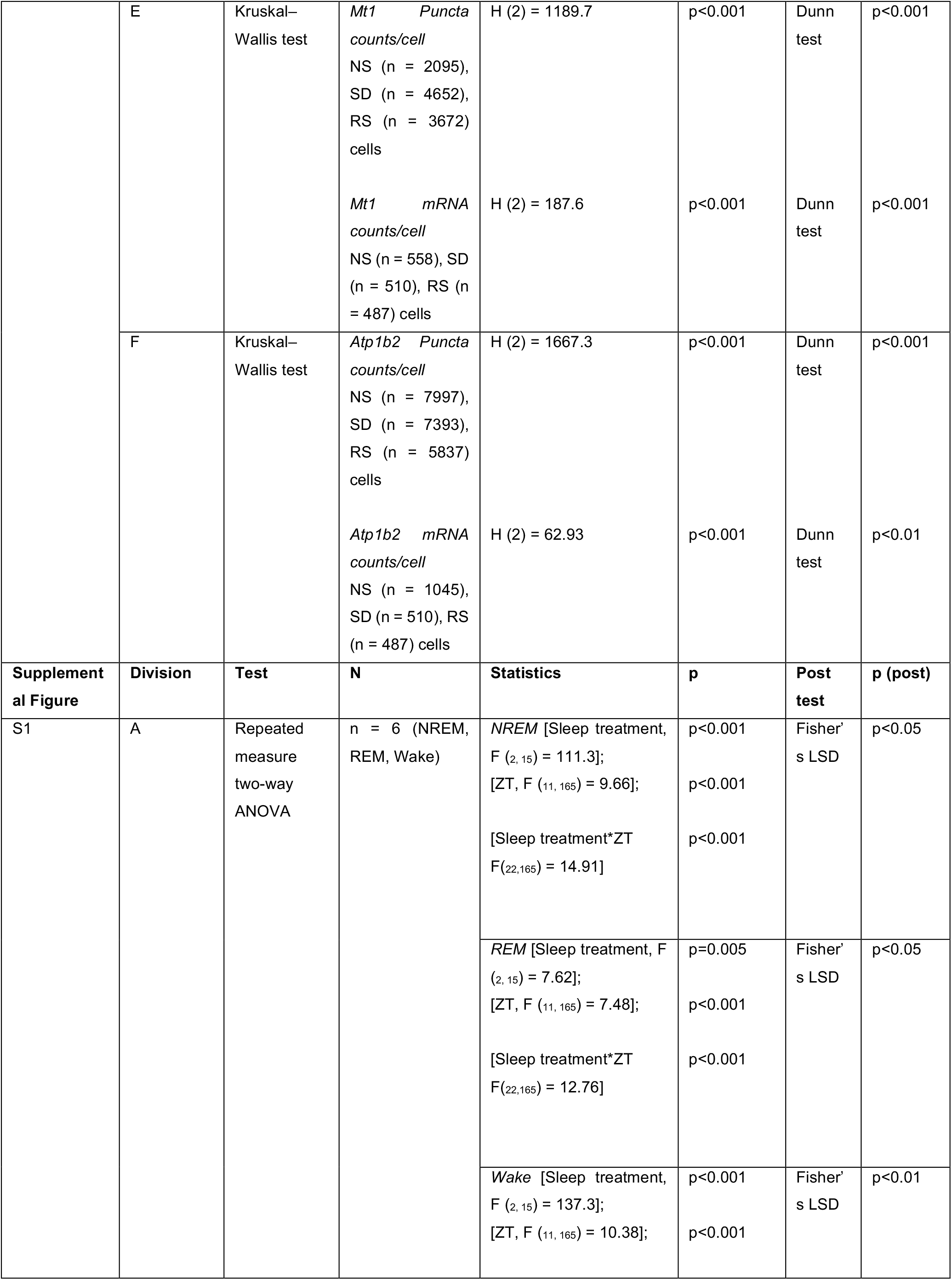
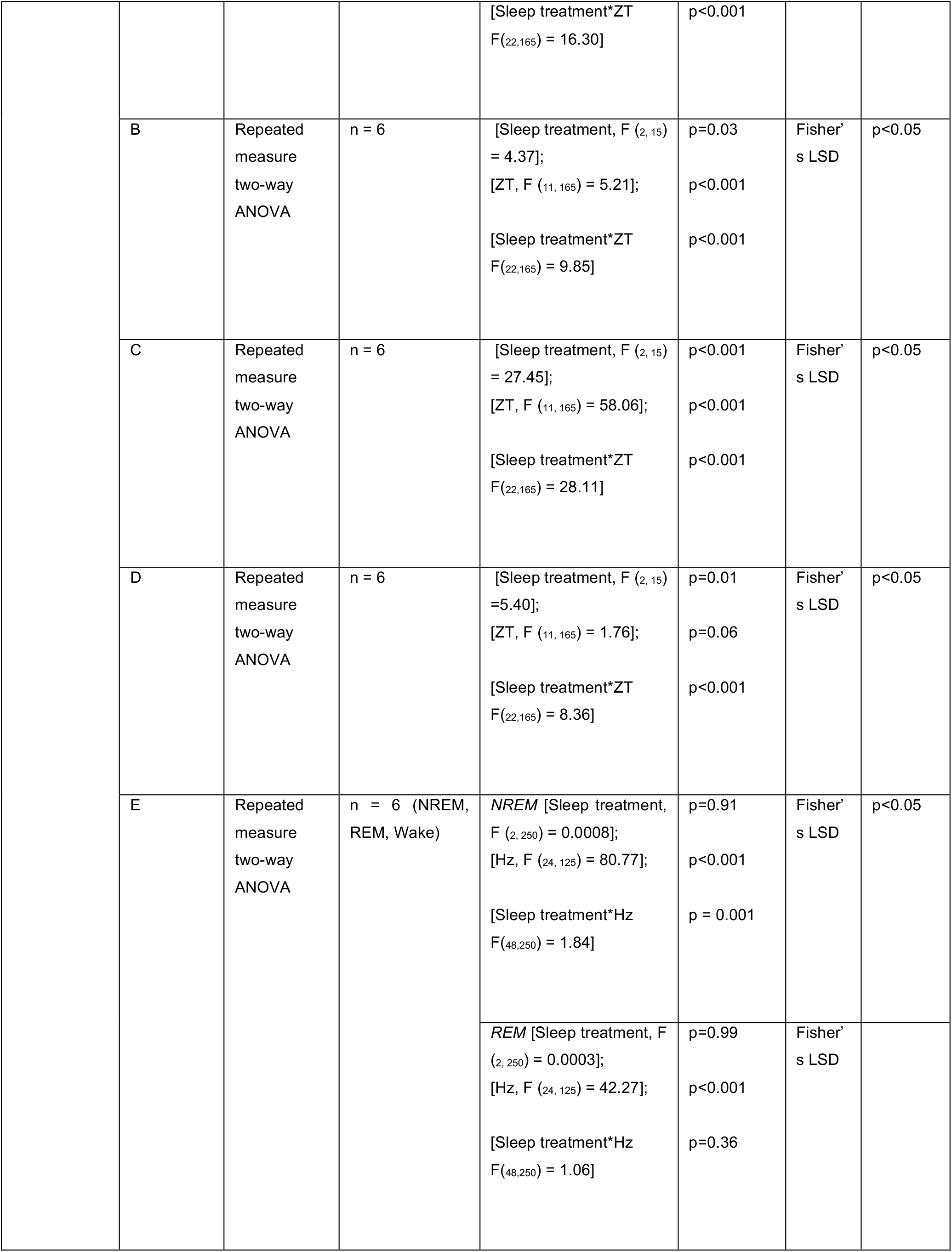
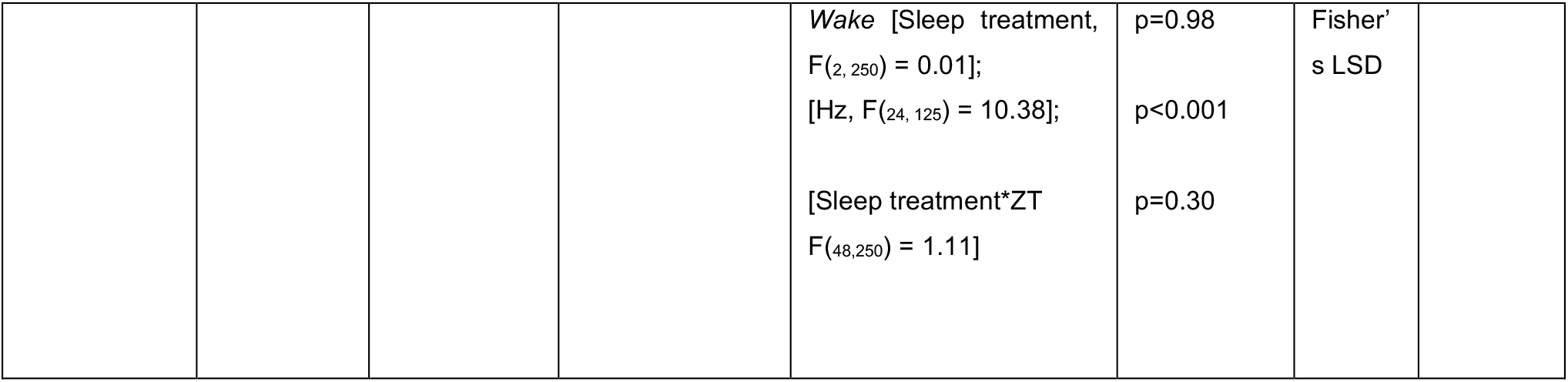
Summary of statistical tests performed.

**Table S2.**
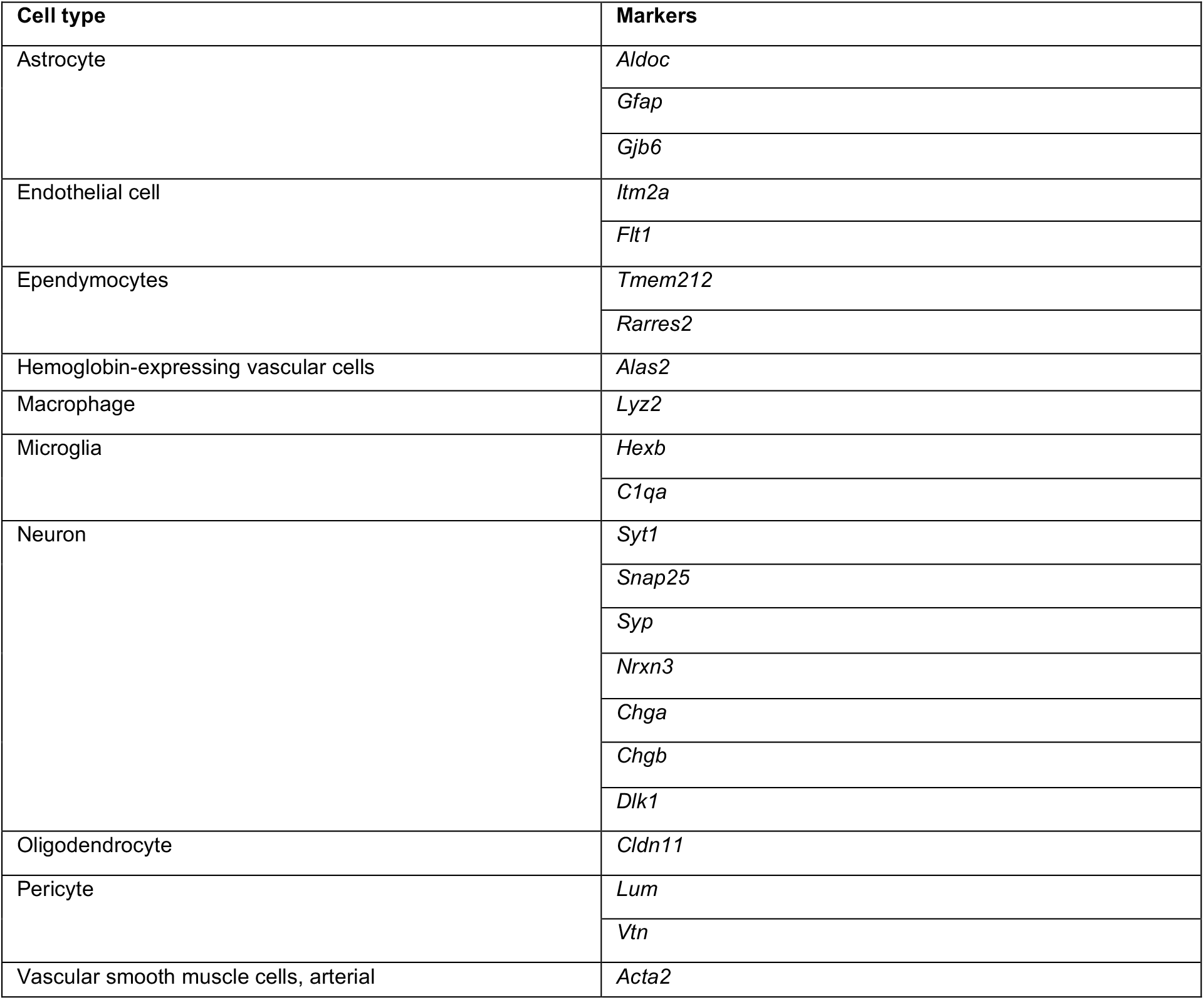
Summary of cell type classifications in single cell transcriptomics.

**Table S3.** Summary of differential gene expression within the cell types of brainstem, cortex and hypothalamus.

| Brain area | Cell type | Significant altered genes after sleep treatments |
| --- | --- | --- |
| Brainstem | Astrocyte | 141 |
|  | Endothelial | 169 |
|  | EPC | 83 |
|  | Microglia | 171 |
|  | Neuron | 84 |
|  | Pericyte | 124 |
|  | VSMCA | 66 |
| Cortex | Astrocyte | 80 |
|  | Endothelial | 92 |
|  | Microglia | 122 |
|  | Neuron | 110 |
| Hypothalamus | Astrocyte | 482 |
|  | Endothelial | 58 |
|  | EPC | 189 |
|  | Microglia | 25 |
|  | Neuron | 220 |
|  | Pericyte | 3 |
|  | VSMCA | 7 |

**Table S4.**
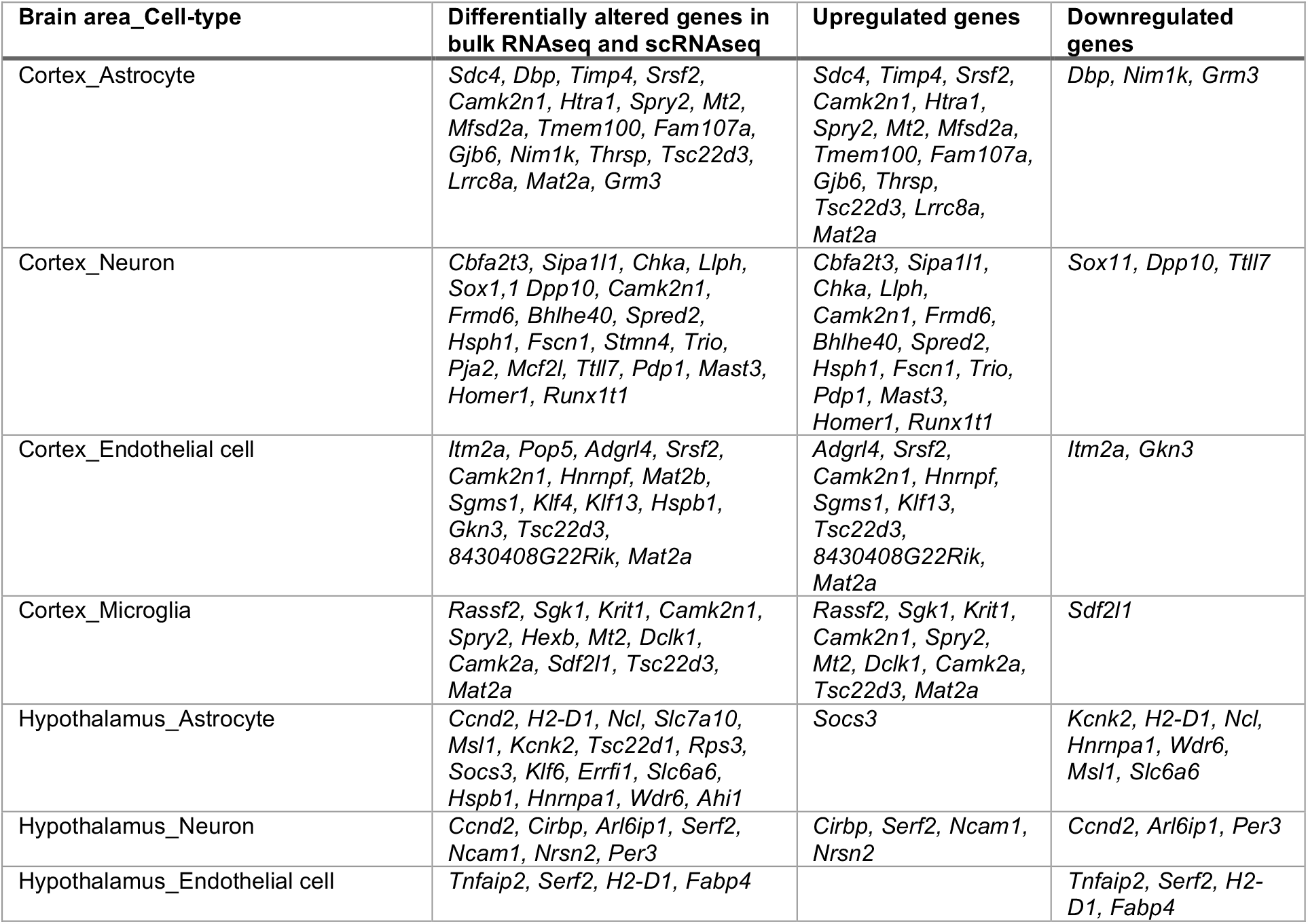
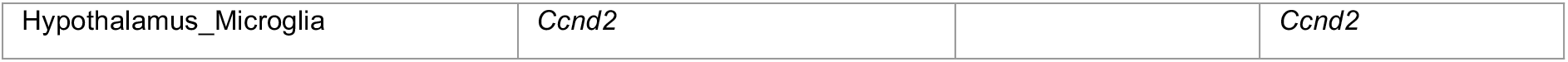
Cell-type specificity of previously reported sleep related upregulated and downregulated genes in the cortex and hypothalamus

**Table S5.**
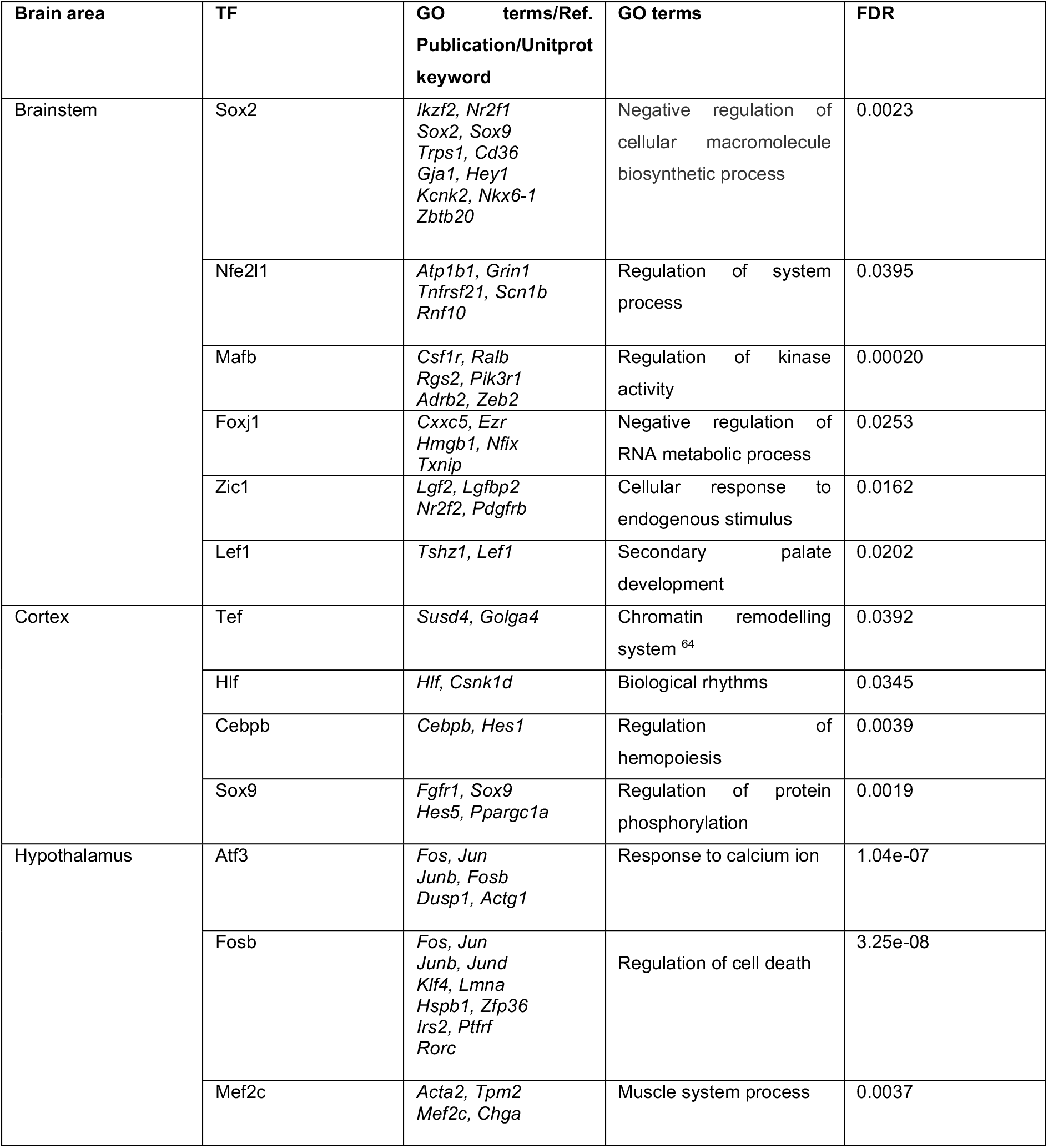
Gene annotation enrichment analysis results for uniquely expressed transcription factors and co-expressed target genes from cells in sleep deprived mice.

**Table S6.** Functional annotations of total and differentially expressed global proteins in astrocytes across sleep treatments.

| Astrocyte (total expressed global proteins) | Astrocytes (differentially expressed global proteins across sleep treatments) |
| --- | --- |
| GOBP:0006091: generation of precursor metabolites and energy | GOBP:0050808: synapse organization |
| GOBP:0050808: synapse organization | GOBP:0061572: actin filament bundle organization |
| GOBP:0043254: regulation of protein complex assembly | GOBP:0043270: positive regulation of ion transport |
| GOBP:0015980: energy derivation by oxidation of organic compounds | GOBP:0050803: regulation of synapse structure or activity |
| GOBP:1990778: protein localization to cell periphery | GOBP:0032535: regulation of cellular component size |
| GOBP:0001505: regulation of neurotransmitter levels | GOBP:1902414: protein localization to cell junction |
| GOBP:0006887: exocytosis | GOBP:0034776: response to histamine |
| GOBP:0045333: cellular respiration |  |
| GOBP:0046034: ATP metabolic process |  |
| GOBP:0031346: positive regulation of cell projection organization |  |
| GOBP:0032535: regulation of cellular component size |  |
| GOBP:0098693: regulation of synaptic vesicle cycle |  |
| GOBP:0032271: regulation of protein polymerization |  |
| GOBP:0010256: endomembrane system organization |  |
| GOBP:0007015: actin filament organization |  |
| GOMF:0003779: actin binding | GOMF:0003779: actin binding |
| GOMF:0050662: coenzyme binding | GOMF:0043177: organic acid binding |
| GOMF:0051082: unfolded protein binding | GOMF:0022836: gated channel activity |
| GOMF:0032549: ribonucleoside binding | GOMF:0005216: ion channel activity |
| GOMF:0003924: GTPase activity | GOMF:0005244: voltage-gated ion channel activity |
| GOMF:0001882: nucleoside binding | GOMF:0022832: voltage-gated channel activity |
| GOMF:0003735: structural constituent of ribosome | GOMF:0015267: channel activity |
| GOMF:0005525: GTP binding | GOMF:0022803: passive transmembrane transporter activity |
| GOMF:0001883: purine nucleoside binding |  |
| GOMF:0003729: mRNA binding |  |
| GOMF:0005516: calmodulin binding |  |
| GOMF:0035254: glutamate receptor binding |  |
| GOMF:0042625: ATPase coupled ion transmembrane transporter activity |  |
| GOMF:0000149: SNARE binding |  |
| GOMF:0051087: chaperone binding |  |
| GOCC:0043209: myelin sheath | GOCC:0032432: actin filament bundle |
| GOCC:0099572: postsynaptic specialization | GOCC:0015629: actin cytoskeleton |
| GOCC:0005743: mitochondrial inner membrane | GOCC:0001725: stress fiber |
| GOCC:0019866: organelle inner membrane | GOCC:0097517: contractile actin filament bundle |
| GOCC:0008021: synaptic vesicle | GOCC:0042641: actomyosin |
| GOCC:0070382: exocytic vesicle | GOCC:0005884: actin filament |
| GOCC:0030133: transport vesicle | GOCC:0099572: postsynaptic specialization |
| GOCC:0015629: actin cytoskeleton |  |
| GOCC:0098798: mitochondrial protein complex |  |
| GOCC:0031252: cell leading edge |  |
| GOCC:0030659: cytoplasmic vesicle membrane |  |
| GOCC:0012506: vesicle membrane |  |
| GOCC:0005938: cell cortex |  |
| GOCC:0030658: transport vesicle membrane |  |
| GOCC:0022626: cytosolic ribosome |  |

**Table S7.** Functional annotations of total and differentially expressed global proteins in neurons across sleep treatments.

| Neurons (total expressed global proteins) | Neurons (differentially expressed global proteins across sleep treatments) |
| --- | --- |
| GOBP:0006091: generation of precursor metabolites and energy | GOBP:0099504: synaptic vesicle cycle |
| GOBP:0043254: regulation of protein complex assembly | GOBP:0099003: vesicle-mediated transport in synapse |
| GOBP:0046034: ATP metabolic process | GOBP:0001505: regulation of neurotransmitter levels |
| GOBP:0008380: RNA splicing | GOBP:0006836: neurotransmitter transport |
| GOBP:0006897: endocytosis | GOBP:0036465: synaptic vesicle recycling |
| GOBP:0007015: actin filament organization | GOBP:0140029: exocytic process |
| GOBP:0006397: mRNA processing | GOBP:0006897: endocytosis |
| GOBP:0015980: energy derivation by oxidation of organic compounds | GOBP:0007269: neurotransmitter secretion |
| GOBP:0045333: cellular respiration | GOBP:0099643: signal release from synapse |
| GOBP:0032271: regulation of protein polymerization | GOBP:0097479: synaptic vesicle localization |
| GOBP:0001505: regulation of neurotransmitter levels | GOBP:0048488: synaptic vesicle endocytosis |
| GOBP:1902903: regulation of supramolecular fiber organization | GOBP:0140238: presynaptic endocytosis |
| GOBP:0051258: protein polymerization | GOBP:0006898: receptor-mediated endocytosis |
| GOBP:0099003: vesicle-mediated transport in synapse | GOBP:0016082: synaptic vesicle priming |
| GOBP:0099504: synaptic vesicle cycle | GOBP:0098693: regulation of synaptic vesicle cycle |
| GOMF:0003779: actin binding | GOMF:0019829: cation-transporting ATPase activity |
| GOMF:0051015: actin filament binding | GOMF:0042625: ATPase coupled ion transmembrane transporter activity |
| GOMF:0003729: mRNA binding | GOMF:0015662: ATPase activity, coupled to transmembrane movement of ions, phosphorylative mechanism |
| GOMF:0005200: structural constituent of cytoskeleton | GOMF:0000149: SNARE binding |
| GOMF:0050662: coenzyme binding | GOMF:0009678: hydrogen-translocating pyrophosphatase activity |
| GOMF:0031072: heat shock protein binding | GOMF:0022853: active ion transmembrane transporter activity |
| GOMF:0003735: structural constituent of ribosome | GOMF:0042626: ATPase activity, coupled to transmembrane movement of substances |
| GOMF:0019001: guanyl nucleotide binding | GOMF:0015399: primary active transmembrane transporter activity |
| GOMF:0032561: guanyl ribonucleotide binding | GOMF:0017075: syntaxin-1 binding |
| GOMF:0001882: nucleoside binding | GOMF:0051117: ATPase binding |
| GOMF:0032550: purine ribonucleoside binding | GOMF:0008022: protein C-terminus binding |
| GOMF:0032549: ribonucleoside binding | GOMF:0015077: monovalent inorganic cation transmembrane transporter activity |
| GOMF:0031625: ubiquitin protein ligase binding | GOMF:0044769: ATPase activity, coupled to transmembrane movement of ions, rotational mechanism |
| GOMF:0001883: purine nucleoside binding | GOMF:0046961: proton-transporting ATPase activity, rotational mechanism |
| GOMF:0044389: ubiquitin-like protein ligase binding | GOMF:0008553: proton-exporting ATPase activity, phosphorylative mechanism |
| GOCC:0043209: myelin sheath | GOCC:0043209: myelin sheath |
| GOCC:0015629: actin cytoskeleton | GOCC:0070382: exocytic vesicle |
| GOCC:0005743: mitochondrial inner membrane | GOCC:0008021: synaptic vesicle |
| GOCC:0019866: organelle inner membrane | GOCC:0030133: transport vesicle |
| GOCC:0014069: postsynaptic density | GOCC:0048786: presynaptic active zone |
| GOCC:0032279: asymmetric synapse | GOCC:0042734: presynaptic membrane |
| GOCC:0098984: neuron to neuron synapse | GOCC:0150034: distal axon |
| GOCC:0031252: cell leading edge | GOCC:0030672: synaptic vesicle membrane |
| GOCC:0099572: postsynaptic specialization | GOCC:0099501: exocytic vesicle membrane |
| GOCC:0005938: cell cortex | GOCC:0012506: vesicle membrane |
| GOCC:0070382: exocytic vesicle | GOCC:0030658: transport vesicle membrane |
| GOCC:0008021: synaptic vesicle | GOCC:0030659: cytoplasmic vesicle membrane |
| GOCC:0022626: cytosolic ribosome | GOCC:0098984: neuron to neuron synapse |
| GOCC:0098800: inner mitochondrial membrane protein complex | GOCC:0030426: growth cone |
| GOCC:0030133: transport vesicle | GOCC:0044306: neuron projection terminus |

**Table S8.** Functional annotations of total and differentially expressed phosphorylated proteins in astrocytes across sleep treatments.

| **Astrocyte (total expressed phosphorylated proteins)** | **Astrocytes (differentially expressed phosphorylated proteins across sleep treatments)** |
| --- | --- |
| GOBP:0050808: synapse organization | GOBP:0008361: regulation of cell size |
| GOBP:0007269: neurotransmitter secretion | GOBP:0032535: regulation of cellular component size |
| GOBP:1902903: regulation of supramolecular fiber organization | GOBP:0050808: synapse organization |
| GOBP:0043254: regulation of protein complex assembly | GOBP:0007269: neurotransmitter secretion |
| GOBP:0007015: actin filament organization | GOBP:0048588: developmental cell growth |
| GOBP:0006887: exocytosis | GOBP:0043254: regulation of protein complex assembly |
| GOBP:1901880: negative regulation of protein depolymerization | GOBP:0060560: developmental growth involved in morphogenesis |
| GOBP:1990778: protein localization to cell periphery | GOBP:0010769: regulation of cell morphogenesis involved in differentiation |
| GOBP:0070507: regulation of microtubule cytoskeleton organization | GOBP:0009187: cyclic nucleotide metabolic process |
| GOBP:0032271: regulation of protein polymerization | GOBP:0007015: actin filament organization |
| GOBP:0050803: regulation of synapse structure or activity | GOBP:0061387: regulation of extent of cell growth |
| GOBP:0043624: cellular protein complex disassembly | GOBP:1990778: protein localization to cell periphery |
| GOBP:0006897: endocytosis | GOBP:0032271: regulation of protein polymerization |
| GOBP:0051656: establishment of organelle localization | GOBP:0007163: establishment or maintenance of cell polarity |
| GOBP:0032535: regulation of cellular component size | GOBP:0043149: stress fiber assembly |
| GOMF:0051015: actin filament binding | GOMF:0005516: calmodulin binding |
| GOMF:0031267: small GTPase binding | GOMF:0003779: actin binding |
| GOMF:0005516: calmodulin binding | GOMF:0042301: phosphate ion binding |
| GOMF:0017016: Ras GTPase binding |  |
| GOMF:0015631: tubulin binding |  |
| GOMF:0019902: phosphatase binding |  |
| GOMF:0005543: phospholipid binding |  |
| GOMF:0003712: transcription coregulator activity |  |
| GOMF:0005096: GTPase activator activity |  |
| GOMF:0008017: microtubule binding |  |
| GOMF:0004674: protein serine/threonine kinase activity |  |
| GOMF:0019903: protein phosphatase binding |  |
| GOMF:0060090: molecular adaptor activity |  |
| GOMF:0008022: protein C-terminus binding |  |
| GOMF:0044325: ion channel binding |  |
| GOCC:0099572: postsynaptic specialization | GOCC:0099572: postsynaptic specialization |
| GOCC:0031252: cell leading edge | GOCC:0098589: membrane region |
| GOCC:0015629: actin cytoskeleton | GOCC:0008021: synaptic vesicle |
| GOCC:0005938: cell cortex | GOCC:0070382: exocytic vesicle |
| GOCC:0099568: cytoplasmic region | GOCC:0031252: cell leading edge |
| GOCC:0005874: microtubule | GOCC:0030426: growth cone |
| GOCC:0008021: synaptic vesicle | GOCC:0015629: actin cytoskeleton |
| GOCC:0070382: exocytic vesicle | GOCC:0030427: site of polarized growth |
| GOCC:0030427: site of polarized growth | GOCC:0030659: cytoplasmic vesicle membrane |
| GOCC:0030426: growth cone | GOCC:0019898: extrinsic component of membrane |
| GOCC:0030863: cortical cytoskeleton | GOCC:0030133: transport vesicle |
| GOCC:0031312: extrinsic component of organelle membrane | GOCC:0012506: vesicle membrane |
| GOCC:0030055: cell-substrate junction | GOCC:0099522: region of cytosol |
| GOCC:0099522: region of cytosol | GOCC:0030118: clathrin coat |
| GOCC:0043209: myelin sheath | GOCC:0099501: exocytic vesicle membrane |

**Table S9.** Functional annotations of total and differentially expressed phosphorylated proteins in neurons across sleep treatments.

| Neuron (total expressed phosphorylated proteins) | Neuron (differentially expressed phosphorylated proteins across sleep treatments) |
| --- | --- |
| GOBP:0032886: regulation of microtubule-based process | GOBP:0019932: second messenger-mediated signaling |
| GOBP:0007015: actin filament organization | GOBP:0097479: synaptic vesicle localization |
| GOBP:0007163: establishment or maintenance of cell polarity | GOBP:0019722: calcium-mediated signaling |
| GOBP:0034329: cell junction assembly | GOBP:0099504: synaptic vesicle cycle |
| GOBP:0010639: negative regulation of organelle organization | GOBP:0099003: vesicle-mediated transport in synapse |
| GOBP:0007264: small GTPase mediated signal transduction | GOBP:0043254: regulation of protein complex assembly |
| GOBP:0051656: establishment of organelle localization | GOBP:0098693: regulation of synaptic vesicle cycle |
| GOBP:0043244: regulation of protein complex disassembly | GOBP:0051656: establishment of organelle localization |
| GOBP:0097479: synaptic vesicle localization | GOBP:0006887: exocytosis |
| GOBP:0099003: vesicle-mediated transport in synapse | GOBP:0007269: neurotransmitter secretion |
| GOBP:0099504: synaptic vesicle cycle | GOBP:0099643: signal release from synapse |
| GOBP:0016358: dendrite development | GOBP:0050808: synapse organization |
| GOBP:0006397: mRNA processing | GOBP:0051258: protein polymerization |
| GOBP:0008088: axo-dendritic transport | GOBP:0070507: regulation of microtubule cytoskeleton organization |
| GOBP:0050808: synapse organization | GOBP:0007044: cell-substrate junction assembly |
| GOMF:0051015: actin filament binding | GOMF:0003779: actin binding |
| GOMF:0017016: Ras GTPase binding |  |
| GOMF:0005516: calmodulin binding |  |
| GOMF:0017048: Rho GTPase binding |  |
| GOMF:0019903: protein phosphatase binding |  |
| GOMF:0005088: Ras guanyl-nucleotide exchange factor activity |  |
| GOMF:0015631: tubulin binding |  |
| GOMF:0005543: phospholipid binding |  |
| GOMF:0019902: phosphatase binding |  |
| GOMF:0005089: Rho guanyl-nucleotide exchange factor activity |  |
| GOMF:0008017: microtubule binding |  |
| GOMF:0035091: phosphatidylinositol binding |  |
| GOMF:0004112: cyclic-nucleotide phosphodiesterase activity |  |
| GOMF:0003712: transcription coregulator activity |  |
| GOMF:0004697: protein kinase C activity |  |
| GOCC:0015629: actin cytoskeleton | GOCC:0014069: postsynaptic density |
| GOCC:0032279: asymmetric synapse | GOCC:0032279: asymmetric synapse |
| GOCC:0099572: postsynaptic specialization | GOCC:0098984: neuron to neuron synapse |
| GOCC:0098984: neuron to neuron synapse | GOCC:0099572: postsynaptic specialization |
| GOCC:0031252: cell leading edge | GOCC:0031252: cell leading edge |
| GOCC:0005911: cell-cell junction | GOCC:0015629: actin cytoskeleton |
| GOCC:0001725: stress fiber | GOCC:0005874: microtubule |
| GOCC:0005874: microtubule | GOCC:0150034: distal axon |
| GOCC:0005938: cell cortex | GOCC:0008021: synaptic vesicle |
| GOCC:0005925: focal adhesion | GOCC:0030133: transport vesicle |
| GOCC:0030863: cortical cytoskeleton | GOCC:0070382: exocytic vesicle |
| GOCC:0150034: distal axon | GOCC:0044306: neuron projection terminus |
| GOCC:0034399: nuclear periphery | GOCC:0098858: actin-based cell projection |
| GOCC:0099568: cytoplasmic region | GOCC:0099522: region of cytosol |
| GOCC:0030427: site of polarized growth | GOCC:0005911: cell-cell junction |

**Table S10.**
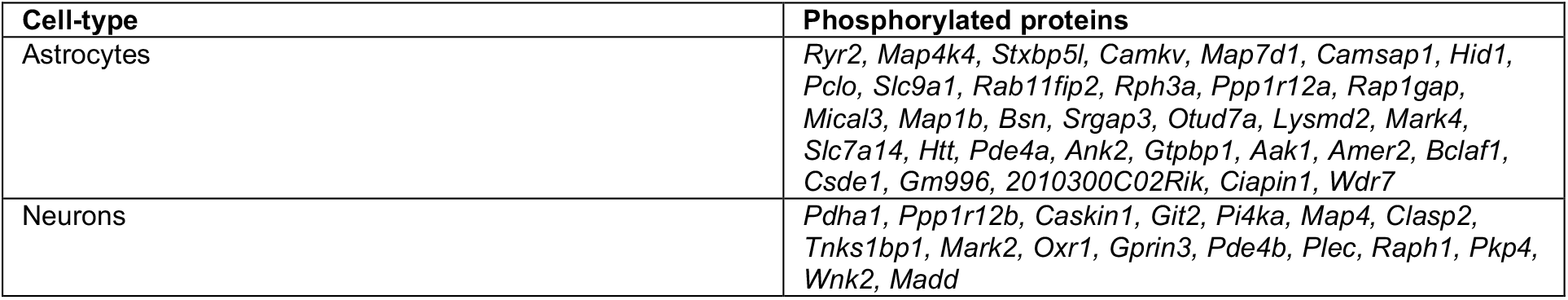
Cell-type specificity to previously known sleep related phosphorylated proteins in cortex

